# BMP signaling promotes heart regeneration via alleviation of replication stress

**DOI:** 10.1101/2024.05.24.595503

**Authors:** Mohan Dalvoy Vasudevarao, Denise Posadas Pena, Michaela Ihle, Chiara Bongiovanni, Pallab Maity, Simone Redaelli, Kathrin Happ, Dominik Geissler, Hossein Falah Mohammadi, Melanie Rall-Scharpf, Chi-Chung Wu, Arica Beisaw, Karin Scharffetter-Kochanek, Gabriele D’Uva, Lisa Wiesmüller, Gilbert Weidinger

## Abstract

One hallmark of aging is a decline in tissue regeneration, which can be caused by DNA replication stress. Whether highly regenerative species like zebrafish are immune from such hindrances to replication is unknown. In contrast to most mammals, adult zebrafish achieve complete heart regeneration via cell cycle entry and proliferation of mature cardiomyocytes. We found that cycling cardiomyocytes experience replication stress, which is induced by the demands of regeneration, but does not occur during physiological heart growth. Since zebrafish cardiomyocyte regeneration is remarkably efficient, heart regeneration appears to depend on elevated capabilities to overcome replication stress. Indeed, pharmacological inhibition of ATM and ATR kinases revealed that DNA damage response signaling is essential for heart regeneration. Using inducible overexpression of ligands and inhibitors of the Bone Morphogenetic Protein (BMP)-Smad pathway, combined with analysis of genetic mutants, we found that BMP signaling alleviates cardiomyocyte replication stress. In the absence of BMP signaling, cardiomyocytes become arrested in the S-phase of the cell cycle, which prevents progression to mitosis and results in heart regeneration failure. Interestingly, BMP signaling can also rescue neonatal mouse cardiomyocytes and human fibroblasts from hydroxyurea-induced replication stress. DNA fiber spreading assays in human cancer cells and human hematopoietic stem and progenitor cells (HSPCs) indicate that BMP signaling acts directly on replication dynamics by accelerating DNA replication fork progression and by facilitating their re-start after replication stress-induced stalling. Our results identify the ability to overcome replication stress as key factor for the elevated heart regeneration capacity in zebrafish. Notably, the conserved capability of BMP signaling to promote stress-free DNA replication might unlock new avenues towards anti-aging and pro-regenerative applications in humans.

## Introduction

Myocardial infarction is a leading cause of death, since the human heart, like that of most adult mammals, lacks the ability to replace cardiomyocytes (CMs) lost to injury. In contrast, adult zebrafish and a few other highly regenerative animals can repair heart injuries via dedifferentiation and proliferation of spared, mature CMs located at the wound border ^1–6^. Zebrafish CM regeneration is remarkably efficient and complete; in response to cryoinjuries that kill 1/3 of the ventricular CMs, pre-injury CM numbers are restored within 30 days ^7^. Many signaling pathways have been found to promote zebrafish CM dedifferentiation and cell cycling ^8–10^. We have previously shown that BMP signaling is required for CM dedifferentiation and proliferation during heart regeneration, but not for CM proliferation during physiological heart growth ^11^. This suggests that BMP signaling primarily acts on a cellular process that is specific to regeneration. Here we propose that BMP signaling promotes CM regeneration by alleviation of replication stress.

Replication stress and other forms of DNA damage are considered leading causes of declining tissue renewal and repair in aged mammals ^12–14^. Cells experience replication stress when DNA replication forks slow down or stall, which can happen due to a variety of reasons, including forks encountering unresolved DNA lesions like DNA strand breaks, collisions with transcriptional machinery or due to insufficient nucleotide supplies ^15,16^. Replication stress can be induced by the high demands on cell cycling imposed by oncogenes or when aged stem cells are challenged to repair a tissue ^16,17^. While cells can use several mechanisms to bypass or tolerate hindrances to replication, unresolved replication stress can result in reduced tissue turnover and repair, functional decline of stem cells, cellular senescence, and apoptosis ^15–19^.

Here, we find that cycling CMs experience replication stress during zebrafish heart regeneration. We identify BMP signaling as a key factor that allows CMs to alleviate the stress, and find that this function is conserved in mammalian cells. Our findings add to the emerging view that highly regenerative organisms are not immune from challenges that impair regeneration in aged mammals. Rather, our work suggests that the ability to efficiently overcome replication stress is a key reason for the elevated capacity of adult zebrafish to regenerate the heart.

## Results

### Cardiomyocytes become γH2a.x positive during zebrafish heart regeneration

To identify mechanisms of heart regeneration, we analyzed the transcriptome of zebrafish hearts, and noticed that gene signatures associated with DNA repair are enriched at 7 days post injury (dpi), which represents the time-point of peak CM proliferation ^1,3–5^ (**Figure 1A, Supplementary data files 1 and 2**). qRT-PCR confirmed that genes involved in several DNA damage response pathways are upregulated at 7 dpi in ventricles that had been subjected to cryoinjury (**Figure 1B**). CMs located at the wound border are key to heart regeneration, since they dedifferentiate and proliferate to replace the damaged myocardium ^2^. Using single-cell sequencing data, we found that some of the upregulated DNA-repair associated genes were more strongly expressed in wound border-zone CMs than in remote CMs at 7 dpi (**Figure 1C**). This prompted us to test whether elevated DNA damage can be detected in wound border CMs, using an antibody against a phosphorylated form of the histone variant H2a.x (γH2a.x), a well-established marker of DNA damage. Western blotting detected elevated levels of γH2a.x in ventricular tissue (from which the wound had been removed to enrich for CMs) at 7 dpi (**Figure 1D**). Immunofluorescence showed that a sizable fraction of CMs at the wound border (within 150 µm of the wound) was positive for γH2a.x at 7 dpi (**Figure 1E**). The temporal profile of γH2a.x-positivity closely resembled the profile for the fraction of CMs that are in the cell cycle, with numbers peaking at 7 dpi and reverting to baseline by 30 dpi (**Figure 1F**). Furthermore, the number of γH2a.x-positive CMs decreased with distance from the wound (**Figure 1G)** in a similar manner as reported for the fraction of cycling CMs ^20^. γH2a.x-positive CMs were also observed in regenerating hearts that had been subjected to resection of the ventricular apex (**Supplementary** Figure 1A). Together, these data suggest that CMs in regenerating hearts experience DNA damage that is independent of the type of injury and unlikely to be caused directly by the insult, as it peaks only at 7 dpi.

**Figure 1.**
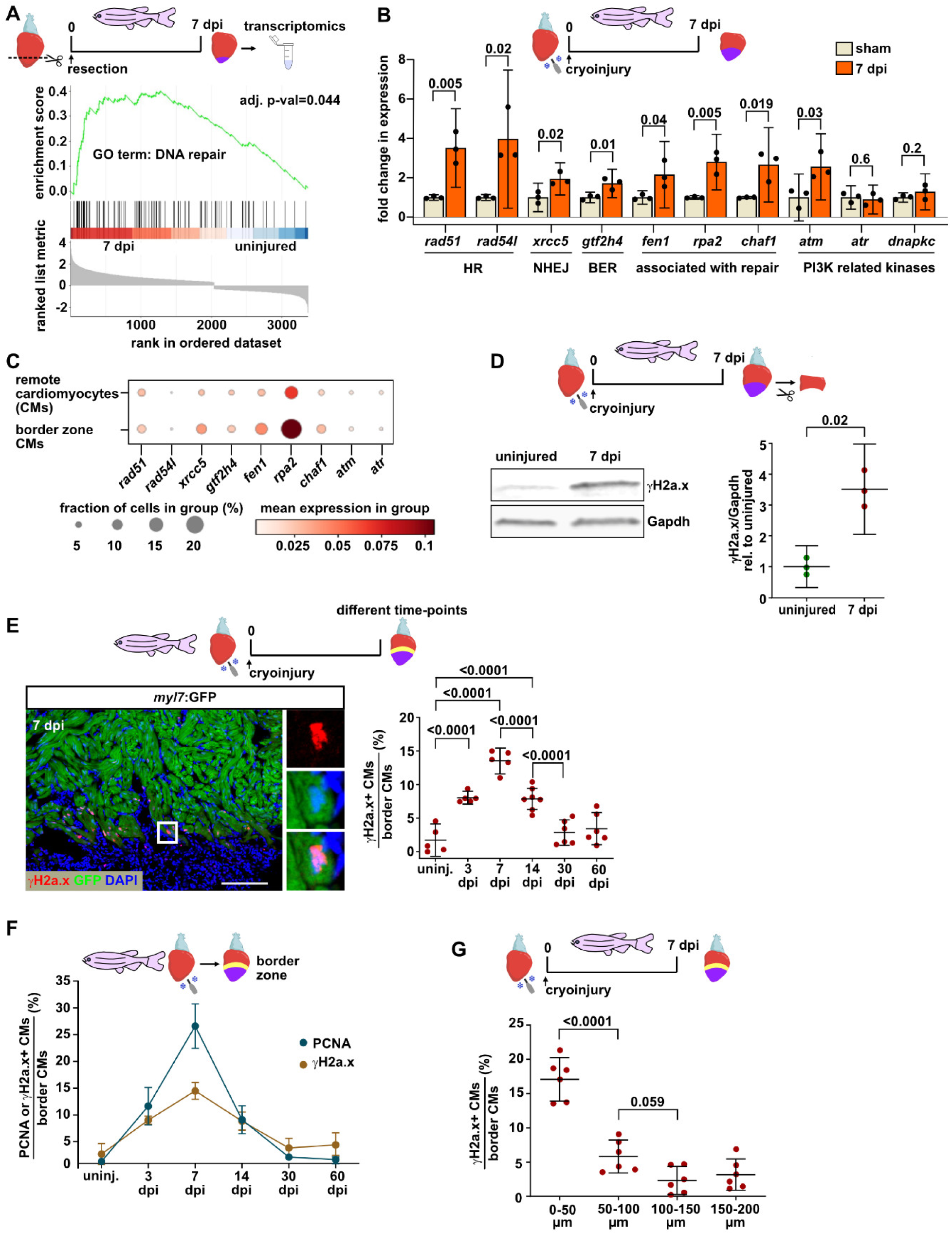
Signatures of DNA damage are induced in regenerating zebrafish cardiomyocytes (CMs) **(A)** Gene Set Enrichment Plot showing enrichment of transcripts associated with the GO term “DNA repair” in zebrafish hearts at 7 days post ventricular resection injury (dpi) compared to uninjured hearts. **(B)** qRT-PCR confirms upregulation of several genes associated with different DNA damage response pathways in zebrafish ventricles at 7 days post cryoinjury (dpi). Expression data is shown relative to the average level in sham treated hearts. Data points represent individual biological replicates, error bars indicate confidence interval (CI) 95%. n_E_ (independent experiments, in this case biological replicates) = 3, n_A_ (animals) = 4 per replicate. Numbers indicate p-value (Student’s t-test). HR, homologous recombination; NHEJ, non-homologous end-joining; BER, base excision repair. **(C)** Dot plot visualization of single cell RNASeq expression data of border zone and remote CM clusters at 7 dpi for the genes upregulated in panel B. **(D)** Western blotting of wound border zone tissue shows increased levels of γH2a.x at 7 dpi relative to uninjured hearts. γH2a.x intensity relative to Gapdh is plotted relative to the average value observed in uninjured hearts. Data points are from independent biological replicates, image shows one representative blot. Error bars, CI 95%; Student’s t-test. n_E_ = 3, n_A_ = 10 per replicate. **(E)** Immunofluorescence on cryosections of *myl7*:GFP transgenic hearts reveals γH2a.x accumulation in GFP+ CMs located within 150 µm from the wound border at 7 dpi. White box in the representative image indicates magnified region. Plot shows fraction in percent of γH2a.x+ CMs out of all CMs located within 150 µm of the wounder border at different time points after cryoinjury. Data points represent fraction observed in individual hearts. Error bars, CI 95%; Analysis of variance (ANOVA) with Bonferroni correction. n_E_ = 1, n_A_ = 5-7 per group, n_C_ (analyzed CMs) = 38200 sum over all time-points. Scale bar, 100μm. **(F)** The fraction of γH2a.x+ wound border CMs correlates with the fraction of cycling CMs. Fraction of γH2a.x+ CMs presented in (E) is overlaid with data of PCNA+ cycling CMs as published in ^7^. **(G)** γH2a.x+ CMs are found close to the wound border at 7 dpi. The fraction of γH2a.x+ CMs is plotted in different bins of distances from the wound border. Error bars, CI 95%; ANOVA with Dunnet’s multiple comparison test. n_E_ = 1, n_A_ = 6, n_C_ = 16400 total.

### Cardiomyocytes experience replication stress during regeneration but not physiological heart growth

The temporal and spatial profiles of γH2a.x accumulation suggested that CMs that enter the cell cycle at the wound border experience DNA damage. To test this, we labeled cycling CMs using repeated injection of EdU and stained for γH2a.x and EdU at 7 dpi (**Figure 2A**). We found that 81 % of the γH2a.x+ CMs were EdU-positive, showing that γH2a.x+ was largely confined to CMs that had recently been cycling or were still in a cycling state. These data suggest that CMs become γH2a.x positive because they experience replication stress when they enter the cell cycle during regeneration. Alternatively, γH2a.x positivity could be a feature of all cycling zebrafish cardiomyocytes. To test this, we labeled cycling CMs in uninjured hearts of juvenile fish (30 days post fertilization [dpf], 16 mm standardized standard length, [SSL]), which undergo physiological growth via CM proliferation, with an 8 h EdU pulse and stained for γH2a.x. We found that only 0.62 % CMs were γH2a.x+, while treatment with hydroxyurea (HU), which depletes cells of nucleotides and thus causes replication stress, did induce γH2a.x+ accumulation (**Supplementary** Figure 2A). Double staining for EdU and γH2a.x revealed that also among the cycling CMs γH2a.x accumulation was rare (2.3 %), but could be strongly increased by HU treatment (**Figure 2B**). Similarly, in adult cryoinjured hearts, HU treatment increased the fraction of γH2a.x+ CMs **(Supplementary** Figure 2B), while it reduced CM cell cycling (**Supplementary** Figure 2C) and the fraction of γH2a.x+ CMs that were also EdU+ (**Supplementary** Figure 2D). Together, these data show that γH2a.x does not generally stain cycling zebrafish cardiomyocytes, but serves as a reliable readout for replication stress. The subnuclear localization of γH2a.x is also indicative of the type of DNA damage ^21^; ionizing irradiation that causes double-strand breaks primarily resulted in accumulation of γH2a.x in distinct puncta in CMs of adult regenerating hearts, while HU caused pan-nuclear γH2a.x staining **(Supplementary** Figure 2E). Importantly, 84 % of the γH2a.x+ CMs of unperturbed regenerating hearts displayed pan-nuclear γH2a.x distribution (**Figure 2C).** Accumulation of phosphorylated Rpa32, which forms part of the RPA-complex of single-stranded DNA binding proteins, is considered a specific readout for replication stress ^22^. Western blotting showed that p-Rpa32 (S33) levels strongly increased by 7 dpi in myocardial ventricular tissue (**Figure 2D**). We conclude that wound border CMs experience replication stress, which can be detected by γH2a.x accumulation.

**Figure 2.**
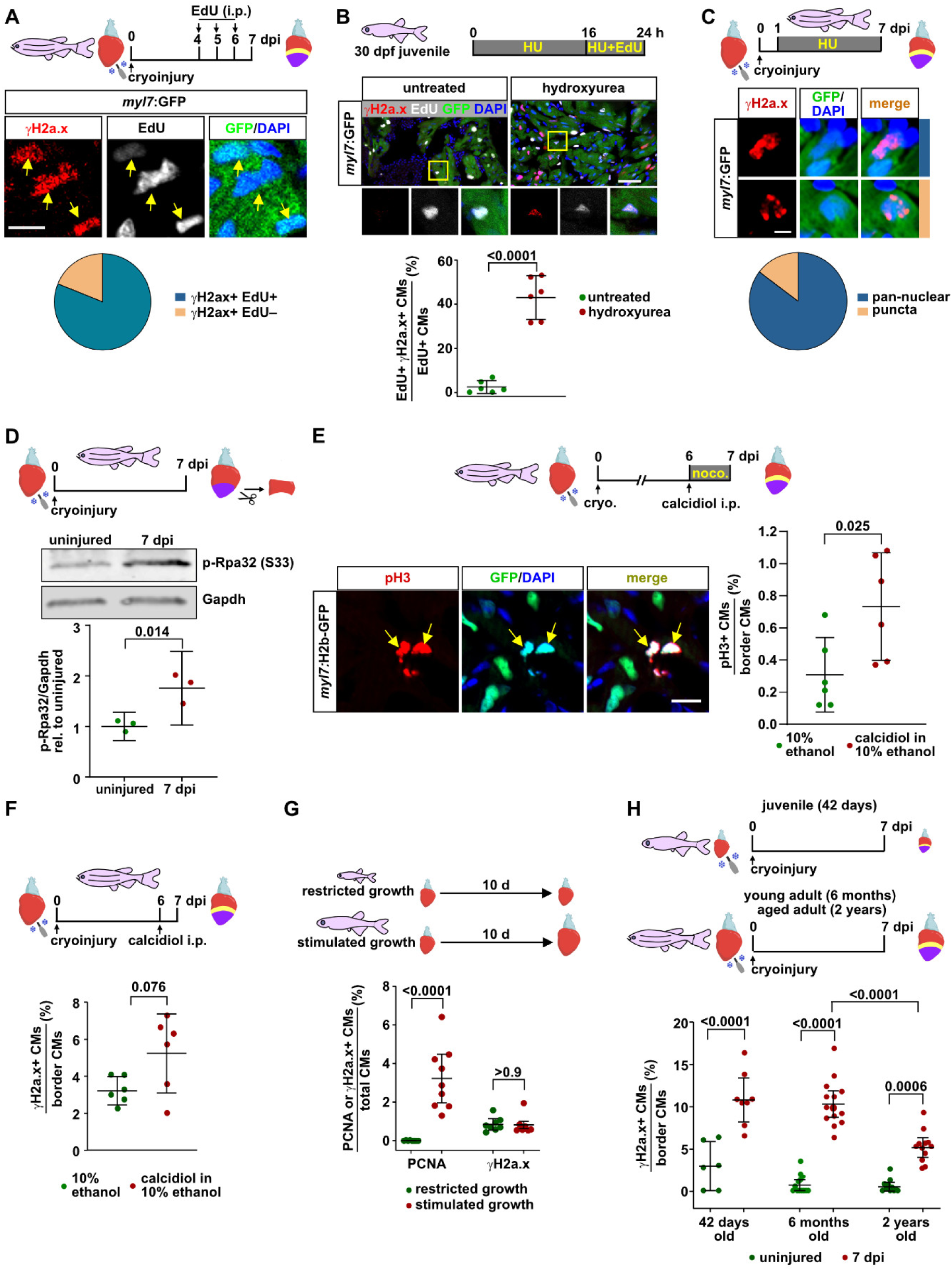
DNA damage upregulation is a specific feature of regeneration and not of physiologic growth or aging. (**A**) Immunofluorescence for γH2a.x and GFP combined with EdU detection in *myl7*:GFP transgenic hearts reveals that γH2a.x is largely confined to GFP+ CMs within the wound border at 7 dpi. Cycling CMs were labeled by daily EdU injections from 4 to 6 dpi. Graph plots how many of the γH2a.x+ CMs at the wound border were EdU+ and EdU–. Error bars, CI 95%. n_E_ = 1, n_A_ = 6, n_C_ = 7170. Scale bar, 10 μm. (**B**) In untreated uninjured juvenile fish at 30 days post fertilization (30 dpf), cycling CMs (labeled by immersion of fish in EdU for 6 hours) are not γH2a.x+, while hydroxyurea treatment induces γH2a.x+ accumulation. Yellow box in the representative image indicates magnified region. Graph plots the fraction in percent of γH2a.x+ EdU+ CMs out of all EdU+ CMs. Error bars, CI 95%, Student’s t-test. n_E_ = 1, n_A_ = 6 per treatment, n_C_ = 4560 untreated, 3660 HU. Scale bar, 50 μm. (**C**) The majority of γH2a.x+ CMs at the wound border at 7 dpi presents pan-nuclear staining (upper image) versus distinct foci (lower image). Error bars, CI 95%. n_E_ = 1, n_A_ = 6, n_C_ = 6450. (**D**) Western blotting of ventricular wound border tissue shows increased levels of Rpa32 phosphorylated at Serin 33 at 7 dpi relative to uninjured hearts. p-Rpa32 intensity relative to Gapdh is plotted relative to the average value observed in uninjured hearts. Data points are from independent biological replicates, image shows one representative blot. Error bars, CI 95%; Student’s t-test. n_E_ = 3, n_A_ = 10 per replicate. (**E**) Immunofluorescence on cryosections of *myl7*:H2b-GFP transgenic hearts reveals an increase in pH3+ mitotic CMs (identified by nuclear GFP) in the wound border area 24 h after a single intra-peritoneal (i.p.) injection of Vitamin D (α-calcidiol) at 6 dpi. Note that nocodazole treatment was used to block cytokinesis, which increases the number of detectable pH3+ cells. Error bars, CI 95%; Student’s t-test. n_E_ = 1, n_A_ = 6 for 10% ethanol, 6 for calcidiol, n_C_ = 6055 for 10% ethanol, 6830 calcidiol. Scale bar, 10 μm. (**F**) Fish injected with α-calcidiol at 6 dpi display a strong trend towards increased fraction of γH2a.x+ wound border CMs at 7 dpi. Error bars, CI 95%, Student’s t-test. n_E_ = 1, n_A_ = 6 per treatment, n_C_ = 5340 EtOH, 4890 calcidiol. (**G**) Stimulated growth condition increases the fraction of PCNA+ CMs without an associated increase in γH2a.x+ CMs in uninjured juvenile fish at 30 dpf (16 mm SSL). Error bars, CI 95%, Student’s t-test. n_E_ = 2 for PCNA and 1 for γH2a.x, n_A_ = 8 restricted growth (RG) PCNA, 8 stimulated growth (SG) PCNA, 8 RG γH2a.x, 7 SG γH2a.x. n_C_ = 83980 RG, 95600 SG. (**H**) The fraction of wound border CMs that are γH2a.x+ at 7 dpi does not increase with fish age. Error bars, CI 95%, Student’s t-test. n_E_ = 1 (42 days), 2 for 6 months and 2 for 2 years, n_A_ = 6 (42 days uninjured), 8 (42 days, 7 dpi), 13 (6 months uninjured), 14 (6 months, 7dpi), 11 (2 years, uninjured), 12 (2 years, 7 dpi), n_C_ = >31700 (total for all conditions).

If heart regeneration induces replication stress, the fraction of γH2a.x+ CMs should be sensitive to interventions that increase or decrease the demand for regenerative CM cell cycling. A single injection of the Vitamin D analog calcidiol, which has been shown to enhance heart regeneration ^23^, increased the fraction of mitotic phospho-Histone H3+ (pH3) wound border CMs at 7 dpi (**Figure 2E**). Interestingly, it also resulted in a strong trend towards an increase in the fraction of γH2a.x+ CMs **(Figure 2F)**. mTOR signaling represents one of the many pathways that have been shown to be required for CM cycling during zebrafish heart regeneration ^24^. We found that treatment with the mTOR inhibitor rapamycin reduced both the fraction of cycling and the fraction of γH2a.x+ CMs at 7 dpi (**Supplementary** Figure 3A). Similarly, starvation yielded comparable results (**Supplementary** Figure 3B). In juvenile fish, the rate of heart growth and thus of physiological CM cycling can be modulated by keeping fish at different densities ^25^. Intriguingly, shifting juvenile fish from restricted to stimulated growth conditions did increase the fraction of cycling CMs, but did not induce γH2a.x+ accumulation (**Figure 2G**). However, γH2a.x+ accumulated in CMs of juvenile fish (42 dpf, 11-16 mm SSL) when their hearts were injured (**Figure 2H**). These findings further support the conclusion that cardiomyocyte replication stress is specifically induced by the demands of regeneration, but not by physiological growth.

In addition to cell cycle entry, wound border CMs undergo dedifferentiation, that is they downregulate characteristics of the differentiated state and re-express embryonic markers ^2^. We thus wondered whether replication stress impinges on CM dedifferentiation. Induction of exogenous replication stress via HU treatment did not affect two readouts of CM dedifferentiation, namely the upregulation of a transgene reporting the activity of regulatory regions of the CM progenitor marker *gata4*, nor the upregulation of an embryonic myosin (embMHC, **Supplementary** Figure 4A). This indicates that CM dedifferentiation occurs independently and likely upstream of CM cell cycling, which is reduced by HU.

### CM replication stress is likely not due to the accumulation of DNA lesions or collisions with transcription

We next sought to probe the molecular reason why regeneration induces replication stress in CMs. One option is that regenerating CMs are particularly sensitive towards blockade of replication forks by DNA lesions that already exist in the DNA prior to S-phase. If so, the fraction of γH2a.x+ CMs might be expected to increase in injured hearts with age of the fish, since lesions should accumulate with age. Yet, the fraction of γH2a.x+ CMs in regenerating hearts at 7 dpi was similar in injured fish at juvenile stages (42 days) and in early-mid adulthood (6 months) and actually decreased in aged conditions (2 years old) (**Figure 2H**). This indicates that CM replication stress is unlikely to be caused by blockade of replication forks by pre-existing DNA lesions. Collision of replication forks with RNA transcription represents another cause of replication stress^16^. Using quantification of antibody staining against the active, elongating phosphorylated form of RNA Polymerase II we surprisingly found that overall levels of transcription appear to be lower in wound border CMs compared to other regions of the ventricle, making it unlikely that proliferating CMs experience increased conflicts between transcription and replication (**Supplementary** Figure 4B).

### DNA damage signaling is required for heart regeneration

We next asked whether CMs that experience replication stress become senescent or undergo apoptosis. We observed the upregulation of the senescence marker beta-galactosidase in the wound area in what appeared by position to be epicardial cells, but not in CMs at 7 dpi (**Supplementary** Figure 4C), in agreement with previous reports in regenerating zebrafish and neonatal mouse hearts ^26,27^. Likewise, while ionizing radiation induced CM apoptosis as detected by Caspase3 accumulation, only 0.27 % of the wound border CMs were apoptotic in unperturbed regenerating hearts at 7 dpi (**Figure 3A**). These results suggest that zebrafish CMs experiencing replication stress are eventually able to overcome this stress and continue to proliferate, despite having retained the capacity to induce pro-apoptotic signaling in response to DNA damage. If so, DNA damage pathways that sense replication stress might be required for heart regeneration. We first tested whether inhibitors of the ATM and ATR kinases, which mediate cellular responses to DNA-damage and replication stress, respectively, are functional in zebrafish. We found that the ATM inhibitor KU55933 and the ATR inhibitor VE821 were non-toxic, since they did not affect embryonic development (**Supplementary** Figure 5A). Yet, embryos treated with the ATR inhibitor displayed necrosis when exposed to doses of HU that on their own had little effect (**Supplementary** Figure 5A), while the ATM inhibitor exacerbated the effect of low doses of ionizing radiation (**Supplementary** Figure 5B). This shows that the inhibitors specifically block the ability of cells to activate repair pathways in response to replication stress and other forms of DNA damage. Intriguingly, treatment of adult fish with both inhibitors in combination impaired morphological heart regeneration, as quantified by shrinkage of the wound between 3 and 21 dpi (**Figure 3B**). Furthermore, treatment for 24 h was also sufficient to impair CM mitosis (**Figure 3C**).

**Figure 3.**
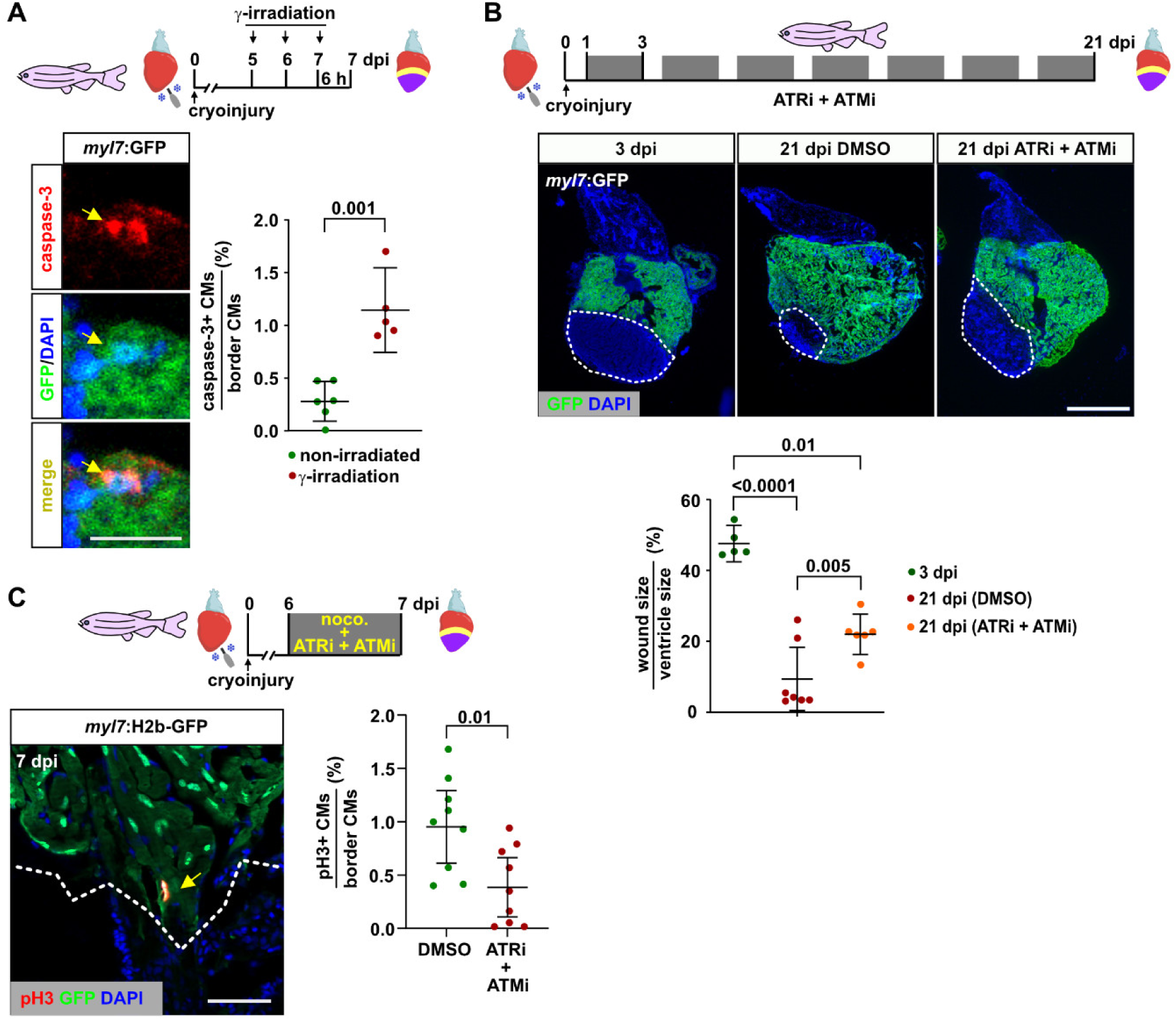
DNA damage repair is essential for zebrafish CM regeneration. (**A**) Very few caspase-3+ apoptotic CMs can be detected at the wound border at 7 dpi, while γ-irradiation induces CM apoptosis. Arrow points to perinuclear caspase-3+ areas in a single apoptotic CM. Error bars, CI 95%, Student’s t-test. n_E_ = 1, n_A_ = 6 for non-irradiated, 5 for γ-irradiation, n_C_ = 3930 non-irradiated, 5950 irradiated. (**B**) Combined intermittent treatment (two days on, one day off) of cryoinjured fish with the DNA damage response inhibitors VE821 (ATR kinase inhibitor) and KU55933 (ATM kinase inhibitor), from 1 dpi to 21 dpi results in an inhibition of wound resorption compared to DMSO treated fish. Wound size (dotted lines) is determined by area lacking GFP expression in *myl7*:GFP transgenics. Error bars, CI 95%, Student’s t-test. n_E_ = 1, n_A_ = 5 for 3 dpi, 8 for 21 dpi DMSO and 7 for 21 dpi ATRi + ATMi. Scale bar, 250 μm. (**C**) Combined treatment with the ATR and ATM inhibitors VE821 and KU55933 for 24 h from 6 to 7 dpi results in a reduction of wound border CM mitosis as determined by anti-pH3 immunofluorescence in *myl7*:H2b-GFP transgenic hearts. Note that nocodazole treatment is used to arrest CMs entering mitosis in a pH3+ state. Error bars, CI 95%, Student’s t-test. n_E_ = 2, n_A_ = 9 per treatment, n_C_ = 16720 DMSO, 15610 ATRi + ATMi. Scale bar, 50 μm.

### BMP signaling alleviates CM replication stress

Our data suggest that alleviation of replication stress is essential for efficient regeneration of CMs to pre-injury numbers in zebrafish. We thus wondered whether specific components of the pro-regenerative signaling environment of the zebrafish heart promote regeneration by impinging on CM replication stress. Likely candidates are pathways that regulate regenerative, but not physiological CM proliferation. We have previously shown that BMP signaling is activated in wound border CMs during zebrafish heart regeneration, but not under conditions of physiological growth, and that it is essential for regenerative, but not physiological CM proliferation ^11^. Therefore, we asked whether BMP signaling modulates CM replication stress. Using a heat-shock-inducible transgene (*hsp70l*:nog3^fr14Tg^), we overexpressed the secreted inhibitor *noggin3* (*nog3*), which interferes with binding of BMP ligands to their receptors and blocks both Smad-dependent and Smad-independent non-canonical BMP signaling (**Figure 4A**). Compared to heat-shocked wild-type fish, *nog3*-expressing fish displayed a significantly higher fraction of γH2a.x+ CMs at 7 dpi (**Figure 4B**). Intriguingly, overexpression of the BMP ligand *bmp2b* (using *hsp70l*:bmp2b^fr13Tg^ fish) very efficiently reduced the number of γH2a.x+ CMs (**Figure 4B**). Importantly, a single pulse of induced *bmp2b* overexpression was sufficient to strongly reduce the number of γH2a.x+ CMs within 6 hours (**Figure 4C**), suggesting that BMP signaling can directly act on cycling CMs that experience replication stress.

**Figure 4.**
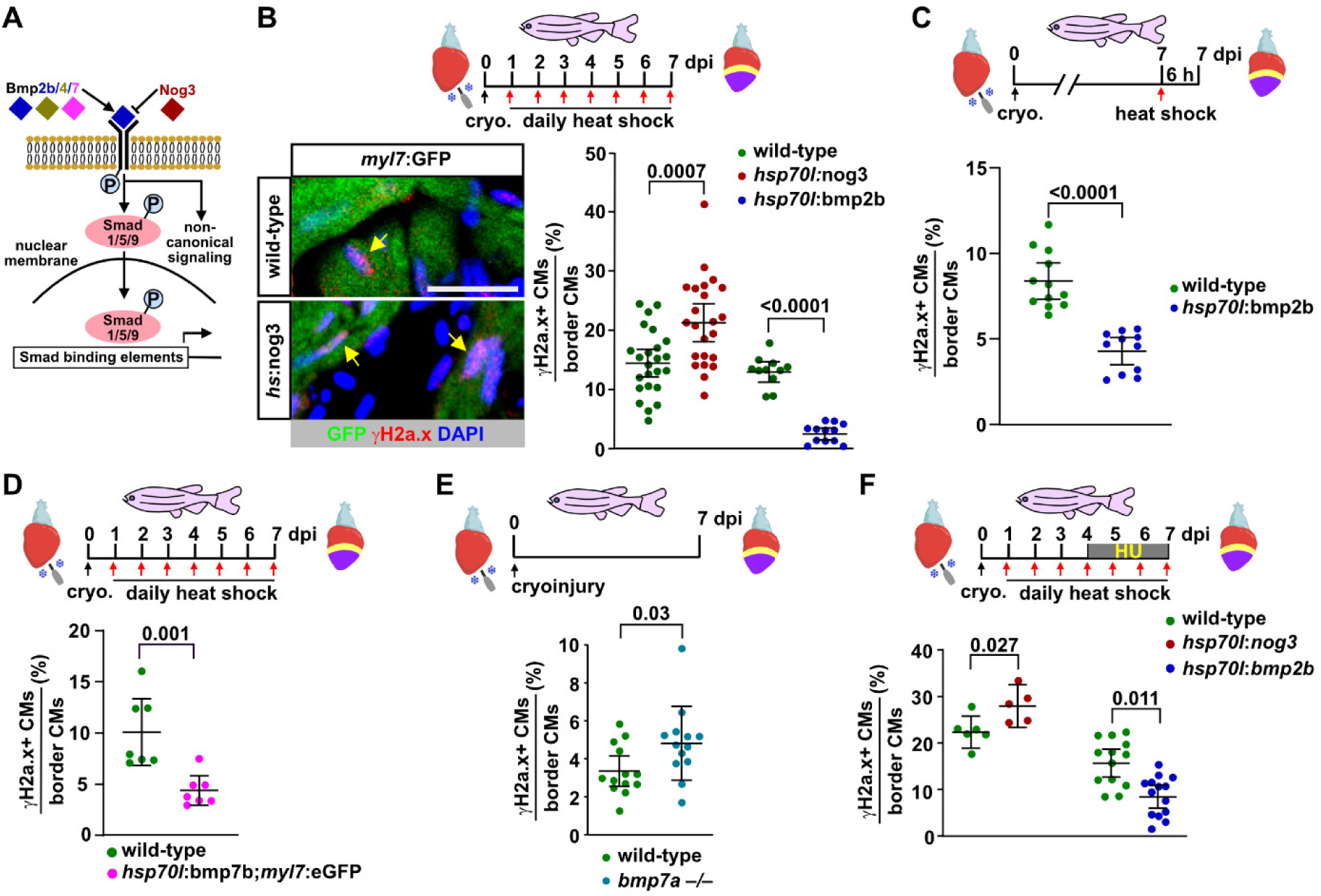
BMP signaling alleviates CM replication stress during zebrafish heart regeneration. (**A**) Cartoon depicting Bmp signaling. Heat shock-induced overexpression of the Bmp ligands 2,4 or 7 from transgenes activates Smad-mediated and “noncanonical” signaling pathways (gain-of-function, GOF), while overexpression of the secreted inhibitor Noggin3 (Nog3) inhibits them (loss-of-function, LOF). (**B**) Bmp-LOF caused by overexpression of *noggin3* via daily heat-shock from 1 to 7 dpi increases the fraction of γH2a.x+ CMs at 7 dpi relative to heat-shocked wild-type siblings, while Bmp-GOF via *bmp2b* overexpression strongly reduces it. Error bars, CI 95%, Student’s t-test. n_E_ = 3, n_A_ = 24 wild-type (wt) siblings to *hsp70l*:nog3, 23 *hsp70l*:nog3, 11 wt siblings to *hsp70l*:bmp2b, 12 *hsp70l*:bmp2b, n_C_ = 41460 total across all groups. Scale bar, 50 μm. (**C**) Bmp-GOF is sufficient to reduce the fraction of γH2a.x+ CMs within 6 h after a single heat-shock of *hsp70l*:bmp2b transgenics at 7 dpi. Error bars, CI 95%, Student’s t-test. n_E_ = 2, n_A_ = 12 wt, 11 *hsp70l*:bmp2b, n_C_ = 4872 wt, 6470 *hsp70l*:bmp2b. (**D**) *bmp7b* overexpression reduces the fraction of γH2a.x+ CMs. Error bars, CI 95%, Student’s t-test. n_E_ = 1, n_A_ = 7 per group, n_C_ = 4900 wt, 3500 *hsp70l*:bmp7b. (**E**) *bmp7a –/–* mutants display an increased fraction of γH2a.x+ CMs at 7 dpi compared to wild-type siblings. Error bars, CI 95%, Student’s t-test. n_E_ = 2, n_A_ = 13 per group, n_C_ = 7800 wt, 7600 *bmp7a –/–*. (**F**) Bmp-LOF using *hsp70l*:nog3 enhances exogenous, hydroxyurea-induced replication stress in border zone CMs, while Bmp-GOF using *hsp70l*:bmp2b transgenics rescues CMs from hydroxyurea-induced γH2a.x accumulation. Error bars, CI 95%, Student’s t-test. n_E_ = 1 for wt vs *hsp70l*:nog3 and 2 for wt vs *hsp70l*:bmp2b, n_A_ = 6 wt siblings to *hsp70l*:nog3, 5 *hsp70l*:nog3, 13 wt siblings to *hsp70l*:bmp2b, 14 *hsp70l*:bmp2b, n_C_ = 47860 total CMs across all groups.

Bmp2 is most closely related to Bmp4, often acts as heterodimer with Bmp7, and homo– and heterodimers of these ligands activate the same set of BMP receptors ^28^. Consistently, overexpression of *bmp7b* or *bmp4* was sufficient to reduce the fraction of γH2a.x+ CMs (**Figure 4D** and **Supplementary** Figure 5C). Heterodimers of Bmp2 or Bmp4 with Bmp7 seem to be the most potent ligands in many contexts, suggesting that Bmp7 is the least redundant of these molecules ^28,29^. Accordingly, we have recently shown that adult fish lacking functional *bmp7a,* one of the two zebrafish *bmp7* paralogs, show reduced numbers of cycling CMs at the wound border ^30^. We thus wondered whether loss of *bmp7a* also affects replication stress during CM regeneration. While *bmp7a* is required for gastrulation, homozygous mutants can be rescued to adulthood by *bmp7a* mRNA injection into embryos ^30,31^. Adult hearts of *bmp7a –*/*–* individuals were indistinguishable by morphology and ventricle size from their wild-type siblings, but did display significantly decreased p-Smad1/5/9 levels at 7 dpi (**Supplementary** Figures 5D and E). Interestingly, the fraction of γH2a.x+ CMs was increased in *bmp7a –*/*–* mutants at 7 dpi (**Figure 4E**). We conclude that endogenous BMP signaling is required for alleviation of CM replication stress during regeneration, and that Bmp7a acts as non-redundant ligand in this process. We also asked whether BMP signaling can alleviate additional, exogenously induced replication stress. Intriguingly, in HU-treated fish, *nog3* overexpression further increased the fraction of γH2a.x+ CMs, while *bmp2b* overexpression strongly reduced it (**Figure 4F**).

### BMPs act through Smads to curb replication stress

BMP ligands can activate Smad-dependent and –independent (non-canonical) signaling pathways (**Figure 5A)**^32^. Since non-canonical BMP-activated pathways have been described to effect the response of mammalian CMs to heart injury ^33^, we asked whether BMPs act through Smads in the alleviation of replication stress in the zebrafish heart. To test this, we created a transgenic line in which the inhibitory Smad6b, together with nuclear Tomato (nT), is expressed after heat-shock (*hsp70l*:nT-p2a-smad6b^ulm16Tg^). Expression of the transgene in adult CMs was mosaic after a single heat-shock, which allowed us to compare nT+ to nT– CMs. Smad signaling as read-out by nuclear accumulation of phosphorylated Smad1/5/9 was suppressed in nT+ CMs within 6 h post heat-shock at 7 dpi (**Figure 5B**). Intriguingly, nT+ CMs were also 3 times more frequently positive for γH2a.x than wild-type or nT– CMs (**Figure 5C**). We conclude that BMP/Smad signaling is required cell-autonomously in CMs for alleviation of replication stress. To test whether overexpressed BMP ligands act solely through the Smad pathway in this context, we analyzed fish double transgenic for *hsp70l*:bmp2b and *hsp70l*:nT-p2a-smad6b. As expected, the fraction of p-Smad1/5/9+ CMs was increased upon *bmp2b* overexpression, and decreased by *smad6*b, but interestingly, in double transgenic fish the fraction of p-Smad1/5/9+ CMs was comparable to that seen in wild-types (**Supplementary** Figure 6A). This allowed us to ask whether co-expression of *smad6b* can reverse the effect of *bmp2b* overexpression on γH2a.x. While fish that only expressed *bmp2b* showed a strong reduction in the number of γH2a.x+ CMs, this effect was completely abrogated by co-expression of *smad6b* (**Figure 5D**). Together, these data indicate that BMP signaling acts directly in CMs and through Smads to alleviate replication stress.

**Figure 5.**
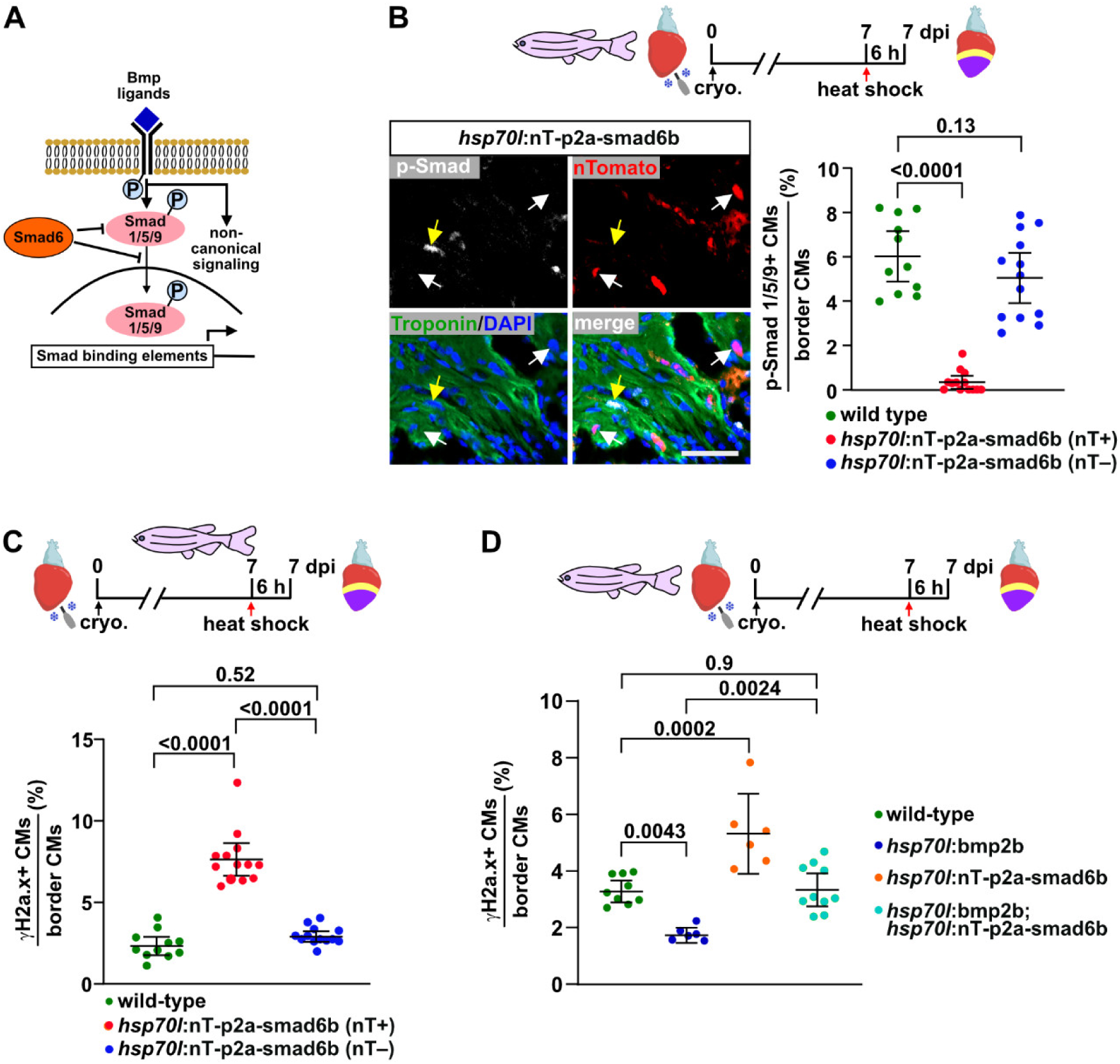
Bmp-mediated alleviation of replication stress requires Smad signaling. (**A**) Cartoon depicting specific inhibition of the canonical Bmp-Smad pathway via overexpression of the inhibitory *smad6b*. (**B**) Immunofluorescence for nuclear Tomato (nT, white arrows) reveals mosaic expression of the *hsp70l*:nT-p2a-smad6b transgene 6 h after a single heat shock at 7 dpi. p-Smad1/5/9 immunostaining reveals inhibition of Bmp-Smad signaling in nT+ CMs (identified by Troponin staining), while nT– CMs in transgenic hearts are equally likely to be p-Smad1/5/9+ than CMs in heat-shocked wild-type fish. Error bars, CI 95%, ANOVA with Bonferroni correction. n_E_ = 2, n_A_ = 11 wt, 13 *hsp70l:*nT-p2a-smad6b, n_C_ = 6400 wt, 4540 nT+, 2950 nT–. Scale bar, 100 μm. (C) Bmp-Smad LOF using *hsp70l*:nT-p2a-smad6b results in an increase in γH2a.x accumulation in nT+ CMs compared with wild-type or nT– CMs after a single heat shock 6 h before harvest at 7 dpi. Error bars, CI 95%, ANOVA with Bonferroni correction. n_E_ =2, n_A_ = 11 wt, 13 *hsp70l:*nT-p2a-smad6b, n_C_ = 5800 wt, 2600 nT+ 2780 nT–. (D) Smad6 co-expression blocks the ability of Bmp2b GOF to alleviate CM replication stress, since the fraction of γH2a.x+ CMs in *hsp70l*:nT-p2a-smad6b; *hsp70l*:bmp2b double transgenics is not different from that in heat-shocked wild-type fish 6 h after a single heat shock at 7 dpi. Error bars, CI 95%, ANOVA with Bonferroni correction. n_E_ =2, n_A_ = 9 wt, 6 *hsp70l*:bmp2b, 6 *hsp70l:*nT-p2a-smad6b, 10 *hsp70l:*nT-p2a-smad6b and *hsp70l*:bmp2b, n_C_ = 13250 total across all groups. The observed relative difference between wild-type and double transgenics is 2%, the calculated smallest significant difference 22%, which is smaller than the difference between *hsp70l*:bmp2b and double transgenics of 47%. Thus, this experiment had enough power to reveal biologically relevant effects and we conclude that Smad6 abrogated the effect of Bmp2b GOF.

### BMP signaling promotes stress-free replication and progression into mitosis

Of note, BMP signaling seems to behave differently than other interventions that alter CM cell cycling and proliferation. We have shown above that the fraction of CMs that cycle and of those that are γH2a.x+ is closely positively correlated for a range of experimental interventions, including inhibition of mTOR and activation of Vitamin D signaling (see Figure 2). In contrast, we found that loss of *bmp7a* results in a decrease in CM cycling ^30^, but an increase in the number of γH2a.x+ CMs (see Figure 4). This prompted us to further explore how BMP signaling regulates regenerative CM cycling and proliferation.

A single pulse of *smad6b* expression reduced the fraction of pH3+ mitotic CMs in cryoinjured *hsp70l*:nT-p2a-smad6b fish within 6 h (**Figure 6A**). This suggests that CMs, which in the absence of BMP signaling cannot overcome replication stress, are stuck in S-or early G2-phase of the cell cycle and cannot proceed into mitosis. To test this idea more directly, we labeled cycling CMs of injured wild type and *hsp70l*:nog3 transgenic hearts for 3 days via repeated EdU injections, and stained for EdU and PCNA at 7 dpi (**Figure 6B**). Our previous quantification and modeling of CM number increase during regeneration has suggested that individual CMs do not undergo multiple rounds of cell division ^7^; thus, a minority of CMs that are cycling at 4, 5 or 6 dpi are expected to still be cycling at 7 dpi, rather many will have withdrawn from the cell cycle by that time. Thus, we considered that EdU+ PCNA+ CMs at 7 dpi are arrested in the cell cycle, while CMs that were only EdU+ had successfully withdrawn from the cell cycle (**Figure 6B**). Indeed, in heat-shocked wild-type fish only ∼35% of the EdU+ CMs were PCNA+ at 7 dpi (**Figure 6B**). Interestingly, inhibition of BMP signaling via overexpression of *nog3* increased the fraction of EdU+ PCNA+ arrested CMs (**Figure 6B**). *smad6b* overexpression increased the fraction of PCNA+ CMs within 6 h after a single pulse of expression (**Supplementary** Figure 6B). Combined with the fact that the same intervention reduced CM mitosis (see Figure 6A), this strongly suggests that *smad6b* also induces CM cell cycle arrest. To test whether CMs were indeed stuck in S-or G2-phase because they could not overcome replication stress in the absence of BMP signaling, we again labeled cycling CMs by repeated EdU injection and then stained for EdU and γH2a.x (**Figure 6C**). We reasoned that EdU+ γH2a.x+ CMs experience replication stress, while CMs that were only EdU+ enjoyed stress-free replication (**Figure 6C**). We found that the fraction of EdU+ γH2a.x+ stressed CMs was increased upon overexpression of *nog3* (**Figure 6C**). Interestingly, also *bmp7a –*/*–* fish displayed a significantly higher fraction of γH2a.x+ EdU+ stressed CMs than wild-type siblings (**Figure 6D**). Overall, these data suggest that, in contrast to other pro-regenerative signaling pathways, BMP signaling specifically promotes CM proliferation because it allows CMs to overcome replication stress and to proceed into mitosis.

**Figure 6.**
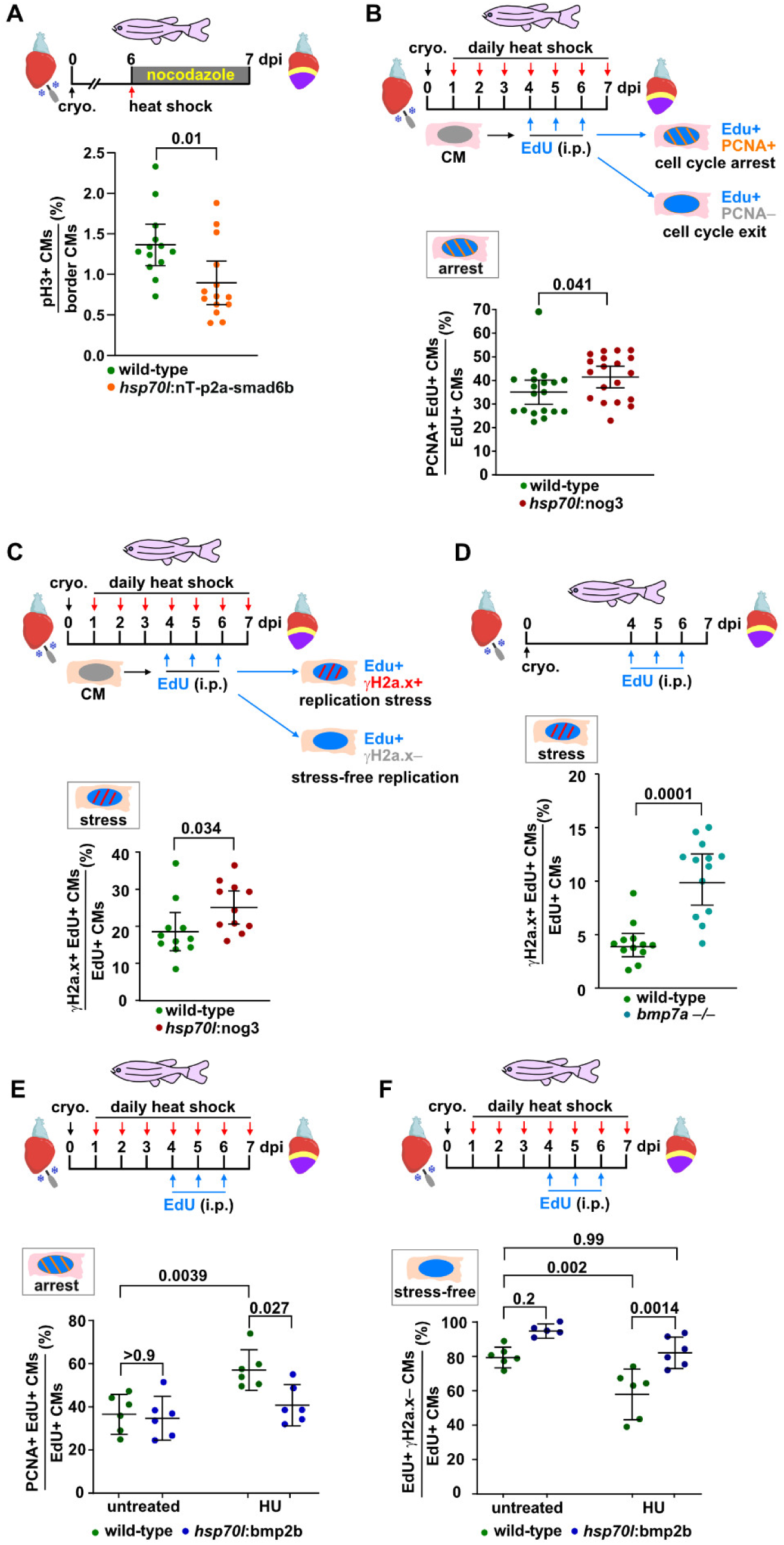
Bmp signaling promotes stress-free replication and relieves CMs from cell cycle arrest. (**A**) Bmp-Smad LOF using *hsp70l*:nT-p2a-smad6b results in a reduction in the accumulation of pH3+ CMs 24 h after a single heat shock administered at 6 dpi. Error bars, CI 95%, Student’s t-test. n_E_ =2, n_A_ = 13 wt, 14 *hsp70l:*nT-p2a-smad6b, n_C_ = 19377 wt, 17328 *hsp70l:*nT-p2a-smad6b. (**B**) Scheme shows how CMs that are arrested in the cell cycle at 7 dpi are distinguished from those that have exited the cell cycle. Cycling CMs are labelled by EdU incorporation from 4 to 6 dpi, followed by EdU and PCNA co-staining at 7 dpi. CMs that are EdU+ PCNA+ display an ongoing cell cycle (most likely due to arrest caused by replication stress). Bmp-LOF using *hsp70l:*nog3 enhances the fraction of EdU+ PCNA+ arrested CMs at 7 dpi over that observed in heat-shocked wild-type fish. Error bars, CI 95%, Student’s t-test. n_E_ =2, n_A_ = 19 wt, 19 *hsp70l:*nog3, n_C_ = 14830 wt, 14981 *hsp70l:*nog3. (**C**) Scheme shows how CMs that experience stress-free replication are distinguished from those experiencing replication stress. Cycling CMs are labelled by EdU incorporation from 4 to 6 dpi, followed by EdU and γH2a.x co-staining at 7 dpi. Double-positive CMs are considered to experience replication stress, Edu+ γH2a.x– CMs are stress-free. Bmp-LOF using *hsp70l:*nog3 enhances the fraction of Edu+γH2a.x– stressed CMs at 7 dpi over that observed in heat-shocked wild-type fish. Error bars, CI 95%, Student’s t-test. n_E_ =1, n_A_ = 11 per group, n_C_ = 5440 wt, 5660 *hsp70l:*nog3. (**D**) *bmp7a –/–* mutants display an increase of the fraction of CMs that experience replication stress at 7 dpi. Error bars, CI 95%, Student’s t-test. n_E_ =2, n_A_ = 12 wt, 13 *bmp7a –/-*, n_C_ = 1500 wt, 1300 *bmp7a –/–*. (**E**) Hydroxyurea-treatment increases the fraction of EdU+ PCNA+ arrested CMs at 7 dpi, which can be reversed by Bmp-GOF using *hsp70l*:bmp2b. Error bars, CI 95%, ANOVA with Bonferroni correction. n_E_ =1, n_A_ = 6 per group, n_C_ = 26000 total CMs across all groups. (**F**) Bmp-GOF using *hsp70l*:bmp2b is sufficient to increase the fraction of CMs that experience stress-free replication at 7 dpi, even when additional stress is induced by HU treatment. Error bars, CI 95%, ANOVA with Bonferroni correction. n_E_ =1, n_A_ = 6 wt (untreated), 5 *hsp70l*:bmp2b (untreated), 6 wt (HU), 6 *hsp70l*:bmp2b (HU). n_C_ = 18910 total CMs across all groups. The observed relative difference between the untreated (wild-type) vs HU treated (*hsp70l*:bmp2b) groups is 12%, the calculated smallest significant difference 14%. Since the smallest significant difference is smaller than the effect size from untreated vs HU treated (wild-type) groups of 20%, we conclude that BMP-GOF can revert HU-induced replication stress down to levels observed in untreated wild-types.

Next, we wondered whether BMP gain-of-function has the opposite effect, namely whether it can promote CM cell cycle progression and stress-free replication. We found that *bmp2b* overexpression was not able to decrease the fraction of arrested PCNA+ EdU+ CMs (**Figure 6E**), possibly because upregulated endogenous BMP signaling in injured hearts is already sufficient. However, induction of additional replication stress via HU treatment increased the fraction of PCNA+ EdU+ arrested CMs in heat-shocked wild-type fish, but *bmp2b* overexpression rescued their numbers back to levels seen without HU treatment (**Figure 6E**). While we observed that *bmp2b* overexpression by itself was insufficient to increase the fraction of EdU+ γH2a.x*–*“stress-free” CMs (**Figure 6F**), it was able to rescue stress-free replication back to normal levels in HU-treated fish (**Figure 6F**).

### BMP signaling protects mouse cardiomyocytes and human fibroblasts from replication stress

We went on to test whether the ability of BMP signaling to alleviate replication stress is conserved in mammals. For a short period of time after birth, mice can regenerate the heart, and CMs isolated from pups at postnatal day 1 (P1) retain proliferative ability in culture ^34^. We have previously shown that BMP7 promotes cell cycle progression and cell division in cultured mouse CMs ^30^. While treatment with either recombinant BMP2 or BMP7 protein did not affect the baseline number of γH2a.x+ CMs in the absence of HU, it abrogated the ability of HU to induce γH2a.x accumulation (**Figure 7A**). We conclude that BMP2 and BMP7 protect neonatal mouse CMs from HU-induced replication stress. Several signaling pathways have been shown to enhance mammalian CM proliferation, including inhibition of p38 kinase activity ^35,36^. We thus wondered whether the ability to abrogate replication stress is a general feature of all pro-proliferative interventions. Yet, the p38 kinase inhibitor SB202190 did not interfere with the ability of HU to induce replication stress (**Figure 7B**). We conclude that the ability to alleviate replication stress is a specific feature of BMP signaling, in agreement with what we have shown above for the regenerating zebrafish heart. Next, we asked whether BMP signaling can protect also other cell types from replication stress. Using primary human neonatal foreskin dermal fibroblasts, we indeed found that combined treatment with recombinant BMP2 and BMP4 proteins completely abrogated the ability of HU to induce γH2a.x (**Figure 7C**). In summary, these data strongly suggest that alleviation of replication stress is a feature of BMP signaling that is conserved over a range of cell types and within vertebrates.

**Figure 7:**
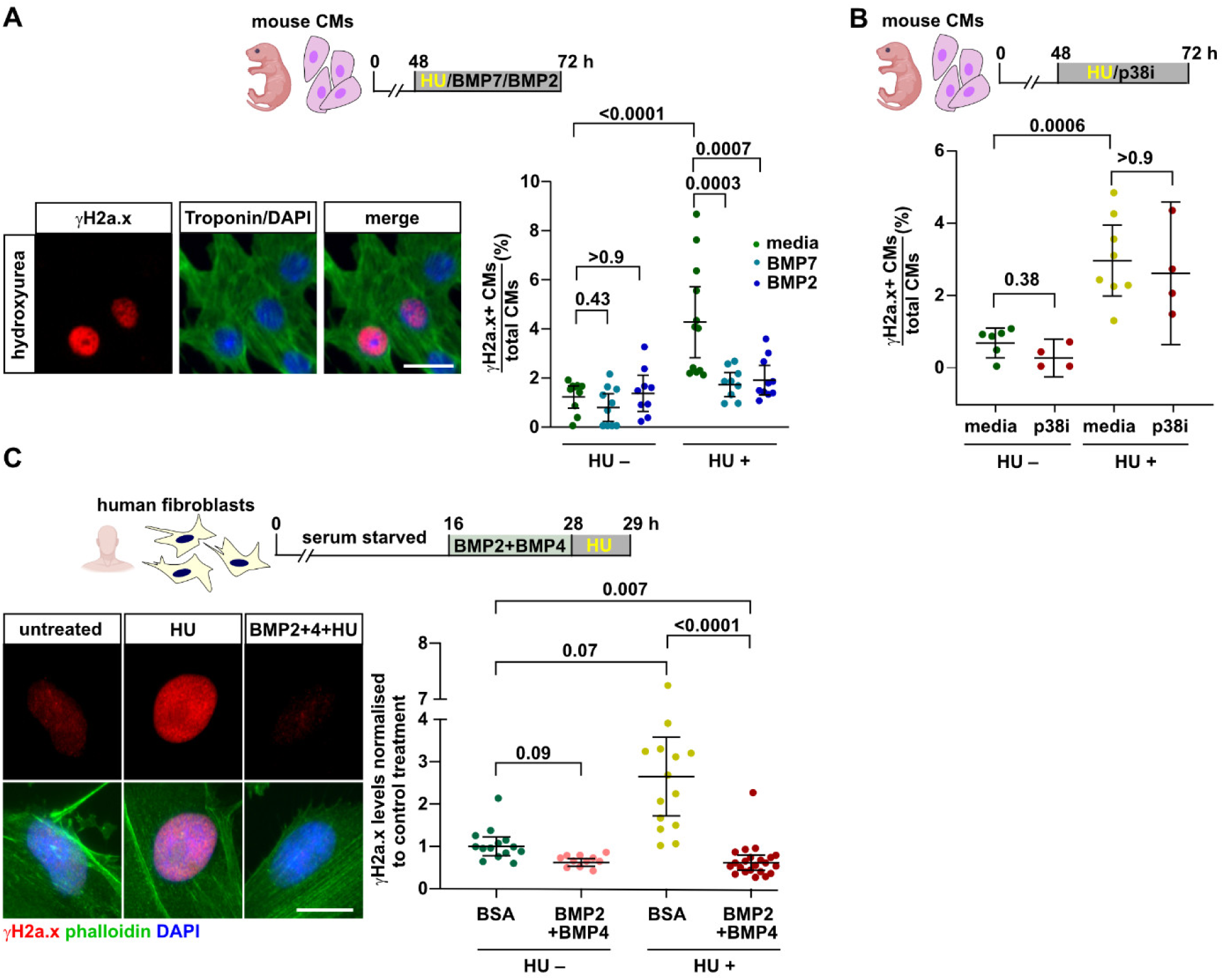
The ability of BMP signaling to alleviate replication stress is conserved in mouse and human cells. (**A**) Immunofluorescence for γH2a.x and Troponin shows that treatment with BMP2 or BMP7 rescues cultured primary neonatal mouse CMs from HU-induced replication stress. Data points represent the fraction of γH2a.x+ CMs per well. Error bars CI 95%, ANOVA with Bonferroni correction. n_E_ =2, n_wells_ = 10 media, 10 BMP7, 9 BMP2, 12 media (HU+), 9 BMP7 (HU+), 10 BMP2 (HU+). n_C_ = 2795 media, 2658 BMP7, 2106 BMP2, 3593 media (HU+), 2704 BMP7 (HU+), 2533 BMP2 (HU+). Scale bar, 20 μm. (**B**) The p38 MAPK inhibitor SB202190, which is known to induce primary neonatal mouse CM cell cycling, does not protect CMs from HU-induced replication stress. Error bars, CI 95%, ANOVA with Bonferroni correction. n_E_ =2, n_wells_ = 6 media, 4 p38i, 8 media (HU), 4 p38i (HU), n_C_ = 1384 media, 765 p38i, 1937 media (HU), 876 p38i (HU). The observed relative difference between HU+ (media) and HU+ (p38i) groups is 11%, the calculated smallest significant difference 36%, which is smaller than the effect size between the HU (media) and HU (BMP7) groups of 44% (from Figure 7A). We conclude that this experiment had enough power to detect effects of that magnitude and that p38i did not rescue CMs from HU-induced replication stress. (**C**) Immunofluorescence for γH2a.x and Phalloidin in human primary neonatal foreskin dermal fibroblasts shows that hydroxyurea induces γH2a.x accumulation, while pre-treatment using a combination of BMP2 and BMP4 prevents it. Data points represent quantification of γH2a.x nuclear levels normalized to the BSA treated control. Error bars, CI 95%, Kruskal-Wallis followed by Dunńs correction. n_E_ =1, n_C_ = 14 BSA, 11 BMP2+4, 14 BSA (HU), 22 BMP2+4 (HU). Scale bar, 20 μm.

### BMP signaling can speed the progression of replication forks and enhances fork re-start

To test whether BMP signaling acts directly on DNA replication, we turned to DNA fiber spreading assays in cultured human cord-blood derived hematopoietic stem and progenitor cells (HSPCs) and the human U2OS cancer cell line. In highly proliferative cells in culture, short incubation with nucleotide analogs (CldU followed by IdU), followed by imaging of individual DNA molecules and quantification of the length of labeled stretches of DNA (“tracks”) can reveal several aspects of replication dynamics, including how fast replication forks progress ^37^. Both cell types encounter replication challenges even under unperturbed culture conditions, HSPCs due to enforced S-phase entry and U2OS cancer cells due to oncogene overexpression ^38–40^. Ongoing replication forks can be identified by dual-color tracks where CldU staining is followed by IdU staining (**Figure 8A**). Intriguingly, treatment with BMP2 or BMP4 recombinant proteins was sufficient to increase the CldU and IdU track lengths of ongoing replication forks in HSPCs **(Figure 8A)**, while in U2OS cells only BMP4 had this effect (**Figure 8B**). Replication fork stalling can be identified as tracks where the CldU+ and IdU+ stretches are of different lengths and can be quantified as the ratio between the longer and shorter stretches ^37^. Except for a very small change caused by BMP2 in HSPCs (long/short track ratio increased by 3.5% from an average ratio of 1.53 in controls to 1.59 in BMP2 treated cells), BMPs did not affect the frequency at which fork stalling occurred in either HSPCs or U2OS cells (**Supplementary** Figure 7A**, B**). Thus, BMP signaling does not appear to be able to prevent fork stalling. Yet, it might alleviate replication stress by promoting re-start of stalled forks. To test this, we first allowed U2OS cells to incorporate CldU into newly synthesized DNA, halted these ongoing replication forks using treatment with HU, and identified re-starting forks after HU-washout and IdU treatment as those where CldU tracks were followed by IdU tracks (**Figure 8C**). Intriguingly, both BMP2 and BMP4 treatment significantly enhanced the fraction of forks that managed to re-start after HU-induced arrest (**Figure 8C**). We conclude that BMP signaling is sufficient to enhance replication dynamics, and propose that it alleviates replication stress at least in part via its ability to enhance replication fork re-start.

**Figure 8:**
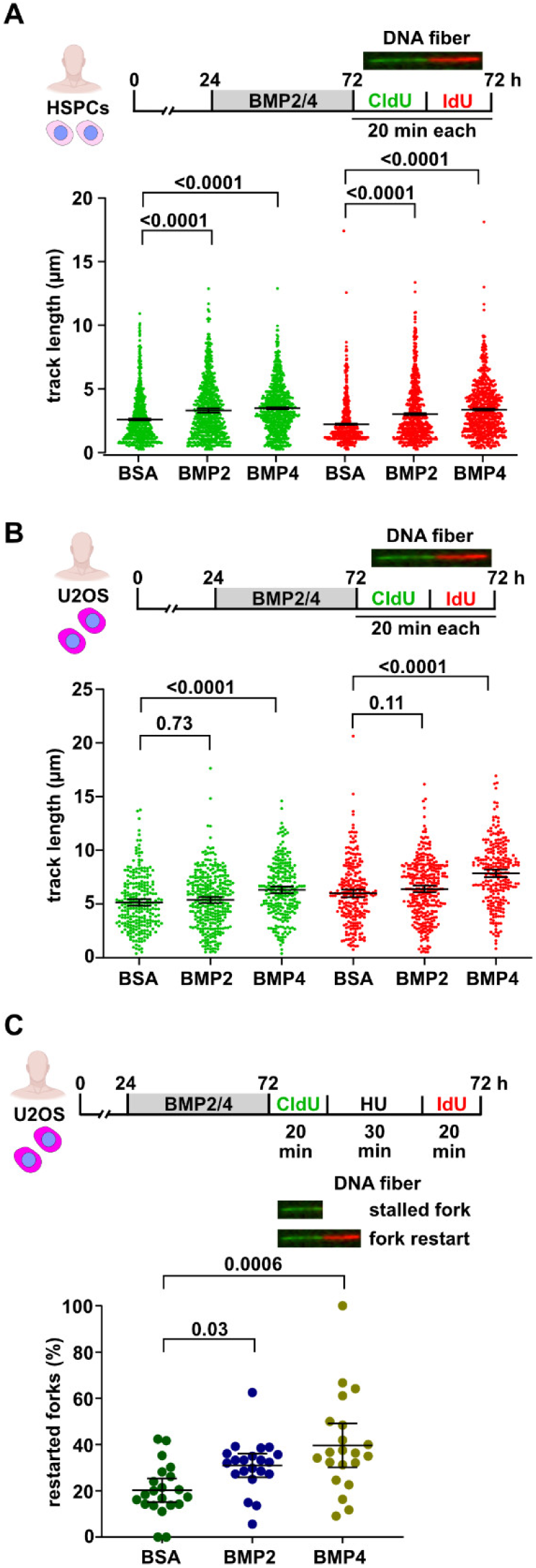
Bmp signaling resolves replication stress by enhancing replication fork speed and re-start. (**A**) DNA fiber spreading assays show that treatment with either BMP2 or BMP4 ligands for 48 h increases the speed of replication fork progression measured by track length of either CldU (green) or IdU (red) in human cord blood hematopoietic stem and progenitor cells (HSPCs). Data points represent length of individual CldU or IdU tracks. Error bars, CI 95%, Kruskal-Wallis followed by Dunńs correction test. n_E_ =3, n_fibers_ > 670 per treatment. (**B**) Treatment with BMP4 increases replication fork progression in the human osteosarcoma cell line U2OS, while BMP2 does not have a significant effect. Error bars, CI 95%, Kruskal-Wallis followed by Dunńs correction. n_E_ =2, n_fibers_ > 250 per treatment. (**C**) Pre-treatment with BMP2 or BMP4 ligands increases the fraction of replication forks that restart after HU-mediated stalling in U2OS cells. Fork arrest is indicated by tracks that are only labeled by CldU, restarted forks by CldU tracks that are followed by IdU. Data points represent fraction of restarted forks out of all analyzed forks. Error bars, CI 95%, Kruskal-Wallis followed by Dunńs correction. n_E_ =2, n_fibers_ > 600 per treatment.

## Discussion

In summary, our data suggest that alleviation of replication stress is a conserved feature of BMP signaling, which is essential for the high capacity of adult fish to regenerate the heart. We propose the following model (**Figure 9**): Cardiomyocyte (CM) regeneration via proliferation of spared CMs is highly efficient and capable of fully restoring pre-injury CM numbers in adult zebrafish. CMs are nevertheless not immune from challenges to DNA replication, which are thought to restrict tissue renewal and stem cell function in aged mammals. In the absence of BMP signaling or upon inhibition of DNA damage response pathways, cycling CMs experiencing replication stress cannot overcome it, get stuck in S-phase of the cell cycle, and regeneration fails. In contrast to other pro-regenerative signaling pathways that promote CM cycling and thereby also increase replication stress, BMP signaling allows for stress-free replication by promoting re-start of stalled replication forks. This ability of BMP signaling is conserved also in mammalian cells; thus, activation of the BMP pathway might have potential as anti-aging and pro-regenerative intervention in aged individuals.

**Figure 9.**
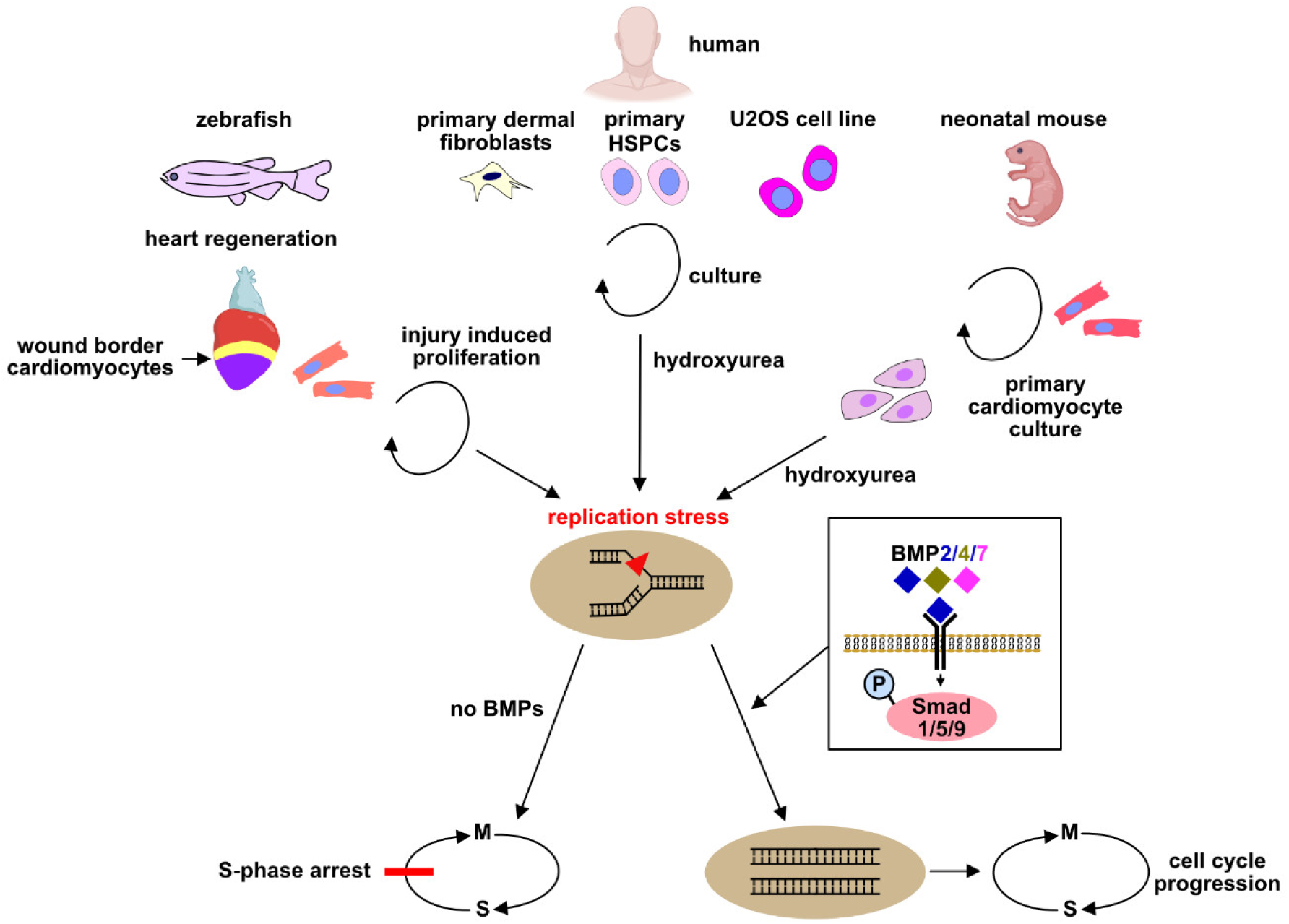
Model for the role of BMP signaling in alleviation of replication stress in zebrafish heart regeneration and mammalian cells. Zebrafish hearts efficiently regenerate via proliferation of pre-existing cardiomyocytes, although cardiomyocytes experience regeneration-induced replication stress. BMP signaling, activated by BMP2, 4 and 7 ligands, and mediated via Smad-signaling allows CMs to overcome the replication stress via promotion of replication fork re-start. This unique ability of BMP signaling to promote stress-free replication is conserved in mammalian cells.

Our findings align with a study by the Whited lab that showed that blastema cells in regenerating axolotl limbs experience replication stress and that interference with DNA damage response pathways impairs limb regeneration ^41^. The fact that highly regenerative systems like the zebrafish heart and the salamander limb are not immune from replication stress appears surprising, but it fits with the realization that efficient regeneration in salamanders can also occur in the presence of cellular senescence, a well accepted confounding factor for tissue renewal and repair in other contexts. In salamanders, limb regeneration can proceed despite the accumulation of senescent cells, since these can be efficiently cleared by macrophages ^42^. In fact, regeneration-induced transient accumulation of senescent cells has been found to be required for regeneration, likely since senescent cells secrete pro-regenerative factors ^43,44^. Of note, our data indicate that zebrafish CMs that experience replication stress do not become senescent. We also do not envision that regeneration-induced replication stress in the zebrafish heart has a beneficial function; rather we assume that it is an unavoidable by-product of the demands that regeneration puts on cell cycling. However, in both the cases of regeneration-induced senescence and regeneration-induced replication stress, successful regeneration appears to depend on efficient mechanisms to overcome these phenomena. We propose that a major reason why BMP signaling promotes CM regeneration in zebrafish is because it allows the cells to alleviate replication stress.

The molecular sources of regeneration-induced replication stress in axolotl blastema cells and zebrafish CMs remain to be identified ^41^, in part due to experimental limitations imposed by the inability to study regeneration *ex vivo*. Possibly, any intervention that results in cell cycle rates exceeding a certain physiological level would cause replication stress, since the high demands on the replication machinery cannot be met. It would be interesting to test whether nucleotides are limiting during regeneration in CMs, which is however challenging to do in intact tissues. Within the limits of sensitivity attainable by the available methods, we found no evidence that collision of replication forks with RNA transcription is likely to cause replication stress. While thousands of transcripts are upregulated during regeneration ^45^, our measurements of RNA-Polymerase II activity did not reveal strongly increased overall transcriptional rates in wound border CMs. Another option is that repair of single-strand DNA breaks does not occur in time before replication forks encounter them. In cancer cells, interventions that push replication fork progression beyond a certain speed have been shown to cause DNA damage because of this reason ^46^.

Alternatively, the high CM cell cycle rates that occur during heart regeneration might not *per se* cause replication stress; rather it might be induced by the specific microenvironment of the injured heart, which differs in several ways from the physiologically growing heart. Candidates could be the immune response, or the production of reactive oxygen species (ROS) ^47^. However, since both ROS and immune responses are required for several aspects of heart regeneration ^47,48^, it would be challenging to test, using available tools, whether they directly modulate CM replication stress. While ROS could play a role, it is worth pointing out that the phenomenon we discovered is clearly distinct from the reported role of oxidative DNA damage in neonatal mouse CMs. Here, the general rise in oxygen tension after birth has been found to cause DNA damage that limits CM proliferation ^49^. Intriguingly, hypoxia was shown to be sufficient to improve CM proliferation in adult injured mouse hearts ^50^. Obviously, there is no reason to assume that oxygen tension would rise in response to heart injury in zebrafish, and thus could be the cause for the accumulation of γH2a.x that we observe. If CMs experience relevant changes in oxygen tension after heart injury, it is likely to be hypoxia, which was shown to decrease DNA damage in mice, yet we do observe an increase in γH2a.x accumulation in the injured zebrafish heart. At a finer scale, an increasing body of evidence shows that regenerating CMs downregulate oxidative phosphorylation and rather switch to glycolysis-dominated catabolism ^51,52^, which again is not compatible with oxidative stress being causative for the observed accumulation of γH2a.x in regenerating CMs. Our data also suggest that γH2a.x accumulation is not due to the fact that cell cycling reveals pre-existing DNA lesions, which could interfere with progression of replication forks, since the fraction of γH2a.x+ CMs does not change with age of the fish. Of note, the oldest fish we analyzed were 2 years old, which is prior to an age (> 3 years) where others have observed declining regenerative capabilities ^53^. Thus, whether functional decline of heart regeneration is associated with increased rates of replication stress is an interesting subject for future research.

We found that BMP-Smad signaling is required for alleviation of CM replication stress in zebrafish regenerating hearts, and that it is sufficient to reduce replication stress in zebrafish and neonatal mouse CMs, as well as in human fibroblasts, U2OS cancer cells and HSPCs. Of note, pre-treatment of cells with BMP ligands can very efficiently protect them from hydroxyurea-induced replication stress. Co-treatment (as in the case of mouse CMs) however also strongly reduces the stress, indicating that BMP signaling does not only increase the resilience of cells towards subsequently induced stress. Rather it likely acts more directly in the alleviation of the stress. Our DNA fiber spreading assays suggest that it might, at least in part, do so via promotion of replication fork re-start.

While a role for BMP signaling in alleviation of replication stress has not been described before, it has been implicated in the response of cells to other forms of DNA damage. Knockdown of the BMPR2 receptor in human pulmonary microvascular endothelial cells results in reduced transcription of Rad51 and increased rates of single and double strand DNA breaks, while BMP9 treatment can protect these cells from accumulation of γH2a.x induced by treatment with the DNA cross-linker mitomycin C^54^. These findings suggest that BMP signaling promotes homologous recombination, which represents one of the mechanisms by which stalled replication forks can re-start ^16^. Thus, these observations are in good agreement with our finding that BMP-signaling can promote fork restart in U2OS cells. Another intriguing study has shown that in cultured cells, in which BMP signaling is active and thus Smad1 is phosphorylated at the SXS motif, genotoxic stress can, via the ATM kinase, result in additional phosphorylation of Smad1 at a different site ^55^. The dually phosphorylated Smad1 was shown to be able to suppress degradation of p53 by Mdm2, resulting in p53 stabilization ^55^. When focusing on p53’s canonical roles, these findings imply that active BMP signaling makes cells more sensitive to react to genotoxic stress by cell cycle exit, apoptosis and senescence^55^. Interestingly, this role of BMP signaling appears to be quite different from how we propose it acts in regenerating zebrafish cardiomyocytes. Since CM apoptosis or senescence is hardly observed in regenerating zebrafish hearts despite strong activation of endogenous BMP signaling, it is unlikely that BMP signaling promotes apoptosis or senescence. On the contrary, we hypothesize that it enhances progression of CMs that experience replication stress through the cell cycle and their mitosis. Furthermore, it has been shown that p53 levels are downregulated during salamander limb and heart regeneration, which appears necessary for efficient proliferation of source cells for regeneration, including cardiomyocytes in the zebrafish heart ^56,57^. Thus, it is unlikely that BMP signaling impinges on genomic stress responses in zebrafish CMs via stabilization of p53. In addition, our DNA fiber spreading assays in cultured human cells indicate that BMP signaling can promote replication fork re-start, which also appears to represent a different function than the p53-dependent cell cycle exit observed by Li and colleagues ^55^. However, more recent works have carved out non-canonical roles of p53 at DNA replication barriers in safeguarding the correct choice of the bypass mechanism depending on the p53 level, cell type and differentiation status ^58–60^. Future work will be necessary to clarify to which extent the different ways in which BMP signaling impinges on genomic stress are cell-type and context-specific.

Accumulation of γH2a.x in CMs of injured zebrafish hearts at 3 dpi has also been reported in a previous study, which however did not investigate its significance in wild-type hearts, but found that telomerase dependent lengthening of telomeres in cardiac cells, including CMs, is essential for zebrafish heart regeneration ^27^. In fish lacking *tert* telomerase activity, CM proliferation was reduced, γH2a.x accumulated in many more CMs, and CM senescence could be detected ^27^. Thus, *tert* activity and BMP signaling as identified by our study represent the only pathways known so far that promote zebrafish heart regeneration by protecting CMs from genomic stress. It will be interesting to test in future research whether additional pro-regenerative signals also act through this mechanism.

The emerging view that highly regenerative species and organs, including the zebrafish heart, are not immune from cellular phenomena like replication stress that limit tissue renewal and repair in aged mammals, but that regeneration rather depends on highly efficient mechanisms to overcome these hindrances, suggests that elucidation of such mechanisms could inform future anti-aging strategies.

## Methods

### Zebrafish husbandry and fish lines

All experiments involving zebrafish were approved by the state of Baden-Württemberg and the animal care representatives of Ulm University. Zebrafish were kept under standard conditions at 26-27°C water temperature and a 14/10 h light/dark cycle. If possible, mixed groups of males and females were used for all experiments. Since significantly different results between males and females were not observed in any of the experiments, data from both sexes are reported together. The following previously published transgenic or mutant fish lines were used: *hsp70l*:nog3^fr14tg61^; *hsp70l*:bmp2b^fr13tg61^; *myl7*:GFP^twu34Tg62^; *14.8gata4*:GFP^ae163^; *myl7*:H2b-GFP^zf521Tg64^; *hsp70l*:bmp7b,*myl7*:eGFP^af5Tg30^; *hsp70l*:bmp4,*myl7*:eGFP^af1Tg65^; *bmp7a*^ty68a/ty68a66^. Transgenic or mutant fish were raised together with their wild-type siblings to adulthood; fish were genotyped and separated only shortly prior to any experiment, and wild-type siblings from the same cross were used as negative controls in all experiments.

### Genotyping

*hsp70l*:bmp2b^fr13tg^ and *hsp70l*:nog3^fr14tg^ fish were genotyped by PCR. Genomic DNA was isolated from fin clips using 50 mM NaOH, followed by incubation at 95°C for 20 min or until tissue was friable. Next, 1/10^th^ volume of 1M Tris-HCl, pH 8.0 was added. The following primers were used (the *hsp70l* forward primer is located in the *hsp70l* promoter and works for both transgenes and is combined with transgene-specific reverse primers: *Hsp70l* (ZDB-GENE-050321-1) GTGGACTGCCTATGTTCATCTTATTTTAGGTCTAC; *bmp2b* (ZDB-GENE-980526-474) ACACCTGACCGAGCAACAGC; *noggin3* (ZDB-GENE-990714-8) GTGGCCAGGAAATACGGTATGTTATCCAT. PCR amplification was conducted according to the program: initial denaturation at 94°C for 3 min, followed by 35 cycles consisting of denaturation at 94°C for 30 secs, annealing at 58°C for 30 secs, extension at 68°C for 30 secs and final extension at 68°C for 2 min.

### Creation of the hsp70l:nT-p2a-Smad6b^ulm16Tg^ transgenic line

The following DNA elements were assembled by PCR– and Gibson-assembly-based cloning methods in a vector containing a single I-SceI meganuclease restriction site: zebrafish *hsp70-4* 1.5kb promoter, nuclear localization signal, dTomato, p2a “ribosome skipping” peptide, zebrafish *smad6b* coding sequence (ZFIN gene ID ZDB-GENE-050419-198), SV40 late polyadenylation site. I-SceI mediated transgene insertion was used to create a stable transgenic line. One subline was selected based on its ability to mimic known BMP loss-of-function phenotypes after heat-shock during gastrulation (dorsalization) and widespread expression after heat-shock in adult hearts.

### Rescue of zebrafish bmp7a (snailhouse) mutants & KASP genotyping

For adult experiments involving *bmp7a* homozygous mutants, embryos from incrosses of *bmp7a (snh)*^ty68a^ heterozygous carriers were injected with 300 pg of *bmp7a* mRNA. After raising them to adulthood, genomic DNA was extracted from fin biopsies, diluted 1:6 in nuclease-free H_2_O and subjected to Kompetitive allele specific PCR (KASP) genotyping. The 2X KASP master mix and KASP assay primer mix were designed by LGC Biosearch Technology from the following *bmp7a* CDS sequence (TACTCTTATGAACCCGCGTACACGACCCCGGGACCCCCGCTGGTGACCCAGCAGGACAGTCGCTTTCTCAGTGATGCCGACATGG**[T/G]**GATGAGCTTTGCGAATACAGGTGAGCGTCTTATGAAATTCACCGCATATCATAATTGTTGTTAGGATGAATCAACAGATTGTTTTTGCTCCATT). The cycling program for KASP PCR involved a hot start activation at 94°C for 15 min, 10 cycles at 94°C for 20 sec and at 61°C for 60 sec, 26 cycles at 94°C for 20 sec and at 55°C for 60 sec, followed by 3 cycles at 94°C for 20 sec and at 57°C for 60 sec. FAM fluorophore (T) corresponds to the wild-type allele and the HEX fluorophore (G) corresponds to the mutated one. The expected genotype distribution is 25% wild-type, 50% heterozygous and 25% homozygous mutants. Adult tanks contained an average of 24% homozygous mutant fish.

### Zebrafish heart cryoinjury, resection and heat-shock-based transgene expression

Ventricular resections were performed as previously described ^1^. Cryoinjuries were performed as previously described ^5^ except that a liquid nitrogen-cooled copper filament of 0.3-mm diameter was used instead of dry ice. For most experiments, heart injuries were performed on zebrafish of 4-6 months of age. For the aging experiments zebrafish of 24 months of age were used. For juvenile experiments, fish of 8-9 weeks of age and 11-16 mm standardized standard length were used. Injured fish were kept at a density of 7 fish per 1.5 liters. Efforts were made to keep fish number equal between experimental groups. In case of death or when equal numbers were unavailable fish from a different, easily identifiable wild-type background were added to equalize the numbers. Heat-shocks were applied by heating water containing fish from 27°C to 37°C within 10 min, and reducing temperature after 1 h back to 27°C within 15 min. For long-term experiments, fish were heat shocked once daily for 7 days; for short-term experiments fish were heat shocked once at 7 days dpi and hearts were harvested 5 h after the end of the heat-shock.

### Zebrafish growth manipulations

Juvenile fish 8-9-week-old (11-16 mm standardized standard length) were either maintained at a high density of 40 fish / liter for a period of 10 days (restricted growth) or a low density of 2 fish / 11 liter (stimulated growth). For starvation experiments, control fish were fed normally (a combination of artemia and solid flakes 2-3 times a day) while starved fish were fed with artemia once daily every second day. Fish were starved starting three days prior to injury throughout the entire duration of the experiment.

### Pharmacological interventions

Detailed information on drugs used is listed in **Supplementary Table 1.** For EdU experiments, adult fish were injected intraperitoneally with 10 µL of 10 mM 5-Ethynyl-2′-deoxyuridine diluted in PBS. Juvenile fish were soaked in EdU (1 mM) dissolved in 50 mL E3 medium (5 mM NaCl, 0.17 mM KCl, 0.33 mM CaCl_2_ x 2H_2_O, 0.33mM, MgSO_4_ x 7H_2_O, 0.2% (w/v) methylene blue, pH 6.5). For detection of mitotic cardiomyocytes, control and experimental fish were soaked with 5 μM nocodazole added to 600 mL of fish facility system water to arrest cells in mitosis. Adult fish were treated with hydroxyurea (20 mM), Rapamycin (1 μM), Atm kinase inhibitor KU55933 (1 μM), Atr inhibitor VE-821 (1 μM) by soaking the fish with respective concentrations in 1L of fish water. DMSO was used to dissolve Rapamycin, ATM and ATR kinase inhibitors. Calcidiol was applied by intraperitoneal injection of 10 µL of 200 µM α-calcidiol dissolved in 10% ethanol.

### Microarray analysis and GSEA protocol

Microarray-based transcriptomics of zebrafish heart regeneration was performed on an Agilent platform with custom designed zebrafish probes **(Supplementary data file 1)**. Ventricular resection was used as injury model, and RNA was isolated from whole ventricles. Uninjured hearts were compared with regenerating hearts at 2, 4 and 7 dpi timepoints. Differentially regulated genes between uninjured and 7 dpi groups were identified using the Bioconductor package Limma (Linear Models for Microarray Data, **Supplementary data file 1**). For these DEGs, GO enrichment analysis was performed using R package clusterProfiler ^67^ against the Genome wide annotation for Zebrafish (R package org.Dr.eg.db, DOI: 10.18129/B9.bioc.org.Dr.eg.db) To find interesting functional terms, the analysis was conducted using the biological process category of the GO, with the following parameters: annotation cutoff of min 10 and max 500 counts, and a p-value cutoff of 0.05 **(Supplementary data file 2).** The terms with FDR adjusted (P.adjust) value < 0.05 were considered as significantly enriched. The enrichment score plots were generated using R package enrichplot (https://bioconductor.org/packages/release/bioc/html/enrichplot.html).

### Mining of published single cell RNA sequencing data

Zebrafish regenerating heart single-cell RNA-seq data was taken from the Gene Expression Omnibus (GEO) database under the accession number GSE251856. Clusters representing CMs were identified based on expression (or enrichment) of *tnnt2a*, *ttn.1*, *ttn.2*, *myl7*, and *pln*. Border zone CMs within the CM cluster were based on the presence/enrichment of the following transcripts ^11,68^: *mustn1b*, *tagln*, *myl6*. Differential gene expression analysis (DEG) was performed using Scanpy.

### qRT-PCR

For qRT-PCR on regenerating zebrafish hearts, RNA was isolated from whole ventricles using the RNeasy for fibrous tissue kit (Qiagen) following the manufacturer’s instructions. cDNA was synthesised using the LunaScript RT SuperMix Kit (New England Biolabs, E3010L) as per manufacturer’s protocol. Reactions containing no reverse transcriptase were used to exclude genomic DNA contamination. qRT-PCR was performed using the Luna Universal qPCR Master Mix (NEB, M3003L) in a BioRad machine. Ct values were normalised to the geometric mean of three housekeeping genes *18S rRNA (*ZDB-RRNAG-180607-2)*, ubb (*ZDB-GENE-050411-10) and *Eef1a1l (*ZDB-GENE-990415-52). Differential expression was calculated using –2^^ΔΔCt^ method ^69^. Primer amplification efficacy was determined on serial 1:4 dilutions of cDNA mix prepared from wounded hearts using the same conditions as those used for experimental samples. Primers were only used if their amplification efficacy exceeded 2. Primer sequences are listed in **Supplementary Table 2.**

### Western blotting of zebrafish heart wound border zones

The wound was identified in extracted whole hearts using stereomicroscopy based on its distinctive color relative to unaffected healthy myocardium. Next, it was removed using fine forceps and scissors. The basal part of the ventricle was removed as well leaving the border zone for homogenization. Border zones from 10 hearts were pooled per sample. Border zone tissues were washed with PBS and manually homogenized in 100 µl SDS lysis buffer (63 mM Tris-HCl, 10% glycerol, 5% 2-mercaptoethanol, 3.5% SDS) using Wheatons glass homogenizers in 1 mL plastic tubes until no particles were seen, followed by centrifugation to clear lysates. SDS-PAGE and immunoblots were performed following standard procedures with a Bio-Rad system. Blotting was conducted onto nitrocellulose membranes, which were incubated overnight with primary antibodies. Information about primary antibodies is listed in **Supplementary Table 3**. After three washes 10 min each with TBS (20mM Tris-HCl, 500 mM NaCl, pH 7.4) Li-cor secondary antibodies were utilized for visualization (**Supplementary Table 4**), and quantification was performed within a linear range of exposure using a Li-COR ODYSSEY Imager and Image Studio Light software. Intensities of p-Rpa32 and γH2a.x protein bands were normalised to Gapdh signals from the same lysate and blot.

### γ-Irradiation

Adult zebrafish were irradiated using a Cs-137 source at 3 Gy / min. Adults were irradiated with 40 Gy. Embryos were irradiated with 8 Gy.

### Tissue processing and immunostaining

Zebrafish hearts were fixed in 4% paraformaldehyde (PFA) in phosphate buffer at RT for 1 h. For γH2a.x stainings hearts were fixed for 2 h. Subsequently, they were washed three times for 10 min in 4% sucrose/phosphate buffer and equilibrated in 30% sucrose/phosphate buffer overnight at 4°C. Hearts were embedded and cryosectioned into 10 μm sections. These sections were evenly distributed onto six serial slides to ensure representation from all ventricular areas on each slide. For immunostainings, slides were washed three times with PEMTx buffer (80 mM Na-PIPES, 5 mM EGTA, 1 mM MgCl_2_ pH 7.4, 0.2% Triton-100) and once with PEMTx + 50 mM NH_4_Cl. Slides were blocked in PEMTx/NGS (10% Normal Goat Serum, 1% DMSO, 89% PEMTx) for 1 h at RT in a humidified chamber. The primary antibodies were applied in PEMTx/NGS at 4°C overnight. For PCNA, antigen retrieval was performed by incubating the slides in pre-heated 10 mM sodium citrate at 95°C in a water bath for 10 min. The primary and secondary antibodies used are listed in **Supplementary Tables 3 and 4** respectively. Secondary antibodies were used at a dilution of 1:1000. Nuclei were shown by DAPI (40,60-diamidino-2-phenylindole) staining. Slides were mounted with Fluorsave (Merck Cat 345789) mounting medium. For EdU detection, cryosections were prepared the same manner as for immunofluorescence staining. Slides were washed twice in 3% BSA/PBS for 5 min, once in 0.5% Triton X-100/PBS for 20 min, and twice in 3% BSA/PBS for 3 min. Next, the slides were incubated in the dark with the reaction cocktail (EdU-Imaging kit, baseclick GmbH) as per manufacturer’s protocol for 30 min. Finally, slides were washed for 3 min in 3% BSA/PBS and continued with the immunofluorescence protocol. For β-Galactosidase staining, Senescence β-Galactosidase Staining Kit #9860 (Cell Signaling) was used as per manufactureŕs protocol. Hearts were fixed with 1X Fixative solution (#11674) for 15 min at RT. Hearts were washed twice with 1X PBS, after which β-Galactosidase Staining solution (1X staining solution #11675, 100X solution A #11676, 100X solution B #11677, and X-gal #11678) was added and incubated overnight at 37°C. Stained hearts were then cryosectioned.

### Image acquisition and analysis

All immunofluorescence images were acquired as single optical planes using 20x magnification (unless otherwise specified) either with Leica Sp5/Sp8 confocal microscopes or with a Zeiss AxioObserver 7 equipped with an Apotome. All image quantifications were performed using ImageJ standard functions. Quantifications of the fraction of EdU, PCNA, γH2a.x, p-Smad positive cardiomyocytes were performed manually on 2 to 3 sections per heart containing the biggest wounds located within 150 µm from the wound border. To define this region of interest, the wound border was identified based on cardiomyocyte staining using ImageJ. The wound was selected using the “free hand selection” tool and added to the Region of Interest (ROI) manager. Next the selection was enlarged 150 µM and added to the ROI manager. Both ROIs were selected and XOR function with overlays was performed. In some cases, especially where pH3 was quantified, the number of cardiomyocytes in the border zone was estimated by multiplying the wound border area (measured using ImageJ) with the average density of cardiomyocytes in three separate regions of interest (size: 100 µm^2^) determined by manual counting of cardiomyocyte nuclei within the regions of interest. For each heart the average value was calculated from the analysis of 2-3 sections per heart. In all other cases, the number of all cardiomyocyte nuclei in the ROI was counted manually for each section separately. For cardiomyocyte mitotic index estimation, the number of pH3+ cardiomyocytes was calculated from the analysis of 8 to 10 sections per heart. For measuring p-Pol2 (S5) intensity, CM nuclei were identified using GFP signal in *myl7*:H2b-GFP transgenic hearts with the ‘Particle Analysi’ tool in Image J. A selection (ROI1) was created for these nuclei after setting a ‘Threshold’ to remove background. The wound border zone was identified as described above and a separate selection was created (ROI2). Using the ‘AND’ function in ImageJ, ROI1 and ROI2 were combined to generate a selection for nuclei of the border zone resulting in ROI3. In the p-Pol2 (S5) channel, background removal was performed and ROI3 was applied. Intensity was measured using ‘Measuré command for individual nuclei.

### Mouse cardiomyocyte cultures, drug treatments and immunofluorescence

Postnatal day 0 or 1 (P0-P1) cardiomyocytes were isolated from wild-type C57BL/6 mouse hearts by mechanical digestion, using a magnetic stirrer, and enzymatic digestion in ADS (5 mM glucose, 106 mM NaCl, 5.3 mM KCl, 20 mM Hepes, 0.8 mM Na_2_HPO4, and 0.4 mM MgSO_4_, pH7.4) with 1 mg/ml pancreatin (Sigma #P1750) and 0.45 mg/ml collagenase A (Roche#10103586001), as previously described in ^70^. Eight to ten hearts were used for each digestion which yielded around 1×10^6^ cells. 35,000 were seeded per well in 96 well-plate. Cardiac cells were then cultured and allowed to adhere for 48 hours in 0.1% gelatine-precoated (Sigma# G9391) plates, with DMEM/F12 (Thermofisher), 1% L-glutamine (Sigma #G7513), 1% sodium pyruvate (Life Technologies #11360-039), 1% non-essential amino acids (Life Technologies #11140-035), 1% penicillin and streptomycin (Gibco#15140122), 5% horse serum (Invitrogen #16050-122) and 10% FBS (Gibco#10082147) and maintained in a humidified atmosphere (5% CO_2_) at 37°C. Thereafter, for replication stress analysis, the medium was replaced with medium containing 5% horse serum without FBS to limit fibroblast growth, supplemented with 0.5 mM Hydroxyurea (Sigma) and the following treatments: BMP7 (R&D #5666-BM), BMP2 (R&D #355-BM), or p38 MAPK inhibitor SB202190 (Sigma #S7067) at 10 µM, for 24 h.

Cardiac cells were fixed with cold 4% paraformaldehyde (PFA) solution for 20 min at RT and then washed three times in PBS and once in 3% BSA (Sigma) in PBS for 5 min. Cells were permeabilized with 0.5% Triton-X100 in PBS for 5 min at RT and a blocking solution (PBS supplemented with 5% BSA (Sigma) and 0.1% Triton-X100) was applied for 1 hour at RT. Then the cells were incubated overnight at 4°C with primary antibodies diluted in PBS, supplemented with 3% BSA and 0.1% Triton-X100. Anti-Troponin T (cTnT) was used as a specific cardiomyocyte marker and anti-phospho Histone H2A.X was used to analyse genomic stress. Following, the samples were washed thrice with PBS and incubated with fluorescent cross-adsorbed secondary antibodies (diluted 1:500 in PBS with 1% BSA (Sigma) and 0.1% Triton-X100 (Sigma)) conjugated to Alexa Fluor™ 555 and 488. After 3 washes in PBS, DAPI (40,60-diamidino-2-phenylindole) was applied for 10 min at RT. Samples were then washed two times in PBS and imaged at 20x magnification.

### Human primary fibroblast cell culture, treatments and immunofluorescence

Human foreskin dermal fibroblasts were cultured in DMEM (Thermo Fisher, Cat No: 11965092) with 10% FCS and 1% Pen/Strep at 37°C and 5% CO_2_. Fibroblasts were cultured up to 70-80% confluency, afterwards, fibroblasts were passaged 1:3 using accutase. For the experiments, fibroblasts were seeded at ∼5000 cells/chamber in poly-D-lysin coated 4-chambered slides (Corning, Cat no: 354577) in 1 ml of DMEM. Following attachment, fibroblasts were serum starved for 16 h and then treated with BMP2 and BMP4 (#355-BM-010 and #314-BP-010, R&D Systems, Minnesota, USA) or BSA at a concentration of 50 ng/mL of each ligand for 12 h. The fibroblasts were then treated with hydroxyurea (Sigma, Cat No: H8627) at a concentration of 100 µM for 1 h. Following treatment, the cells were washed with PBS 3 times, fixed in 4% PFA for 10 min at RT, and permeabilized in 0.2% Triton X-100 in PBS for 10 min at RT. Immunofluorescence was performed by blocking with 5% BSA in PBS for 1 h at RT, overnight incubation at 4°C with anti-γH2AX antibody (Millipore, Cat No: 05-6361) diluted 1:200 in Dako antibody diluent (Agilent, Cat No: S0809), three washes with PBS at RT, incubation with 1:400 Alexa flour 555 labeled secondary antibody for 1 h at RT in the dark, three washes in PBS, and staining with phalloidin-Alexa flour 488 for 30 min. After 3 washes with PBS, nuclei were counterstained with DAPI (40,60-diamidino-2-phenylindole) for 5 min at RT. After 3 washes with PBS, slides were mounted in Dako Fluorescence Mounting Medium (Agilent, Cat No: GM304) and imaged at 40x magnification.

### Human U2OS and HSPC cell culture and DNA fiber spreading assays

Human CD34+ hematopoietic stem and progenitor cells (HSPCs) were isolated from cord blood samples with consent of the parents (approval #155/13 from the Ethical Board of Ulm University) and cultured as described in ^39^ using the following procedure: Cord blood samples were diluted 1:2 in PBS and layered on Ficoll® Paque Plus (Sigma-Aldrich by Merck, Darmstadt, Germany) to isolate mononuclear cells. The cell layer was then transferred to a new tube and washed with PBS containing 2% FBS. Red blood cells were removed by incubation in red blood cell lysis buffer (0.15 M ammonium chloride, 10 mM potassium-bicarbonate, 0.1 mM EDTA in water, pH 7.2 – 7.4), followed by another wash in PBS. CD34+ HSPCs were enriched by using the CD34 Micro Bead Kit (Miltenyi Biotec, Bergisch Gladbach, Germany) according to the manufactureŕs description. HSPCs were cultured for 24 h at 37°C, 5% CO_2_ and ambient O_2_ in serum free StemSpanTM SFEM medium (STEMCELL Technologies, Cologne, Germany) supplemented with 1% Penicillin/Streptomycin (Gibco by Thermo Fisher Scientific, Waltham, Massachusetts, USA) and 1% StemSpanTM CC100 (STEMCELL Technologies, Cologne, Germany) before treatment with either 50 ng/ml BMP2 or BMP4 (#355-BM-010 and #314-BP-010, R&D Systems, Minnesota, USA) solved in 0.1% BSA (Sigma-Aldrich by Merck, Darmstadt, Germany) for 48 h.

U2OS cells were cultivated in DMEM high glucose (Gibco by Thermo Fisher Scientific, Waltham, Massachusetts, USA) supplemented with 10% FBS (Biochrom by Merck, Darmstadt, Germany) at 37°C, 5% CO_2_ and ambient O_2_. For BMP stimulation cells were transferred into serum-free medium X-VIVO 10 (Lonza, Walkersville, Maryland, USA). After 24 h in the serum-free medium treatment with BMP ligands was performed for 48 h as described above for HSPCs.

The DNA fiber spreading assay was performed as described before ^58–60^. In brief, cells were labeled with 20 mM 5-chloro-2-deoxyuridine (CldU, Sigma-Aldrich by Merck, Darmstadt, Germany) for 20 min, then washed with pre-warmed PBS, followed by a second labelling with 200 mM 5-iodo-2-deoxyuridine (IdU, Sigma-Aldrich). For HU treatment, cells were incubated after the first labelling with 5 mM HU for 30 min, then washed again with PBS before the second labelling. Cells were then washed, harvested and resuspended. In case of U2OS, cells were subsequently trypsinized and resuspended. Then 2500 cells were spotted on a slide and lysed with 6 µl lysis buffer (0.5% SDS, 100 mM Tris–HCl (pH 7.4), 50mM EDTA) for 6 min at RT. Slides were then tilted ∼20° to allow the spreading of DNA via gravity. After further 6 min drying, DNA was fixed on the slides by incubation for 5 min with methanol-acetic acid solution (3:1). Slides were either dried for another 7 min and stored in 70% (v*/*v) ethanol at 4°C overnight or directly subjected to denaturation*/*deproteination in 2.5M HCl for 1h, followed by immunofluorescence staining.

Blocking with 5% (w*/*v) bovine serum albumin in PBS for 45 min at 37°C was followed by incubation with a primary antibody mix of anti-BrdU detecting IdU (mouse, #347580, BD, Franklin Lakes, NJ, USA) and anti-BrdU detecting CldU (rat, monoclonal, clone BU1/75 (ICR1), BioRad, Cat-Nr: OBT0030) for 1 h at RT. Finally, slides were incubated with a mix of secondary antibodies, AlexaFluor555 (anti-mouse) and AlexaFluor488 (anti-rat, both secondary antibodies were from Invitrogen by Thermo Fisher Scientific, Waltham, MA, USA) for 45 min at RT. DNA fibers were visualized with a BZ-9000 microscope (Keyence, Cologne, Germany) and a 40× objective. Measurement of the DNA fiber track length was carried out with the Fiji software (ImageJ Wiki, Laboratory for Optical and Computational Instrumentation, University of Wisconsin-Madison, WI, USA).

### Statistics

Statistical analyses were performed using GraphPad Prism 9 software. Information about samples sizes and types of statistical tests used can be found in figure legends. Error bars represent CI 95%. If not stated differently, sample sizes are given in the figure legends as follows: n_E_ = number of independent experiments or biological replicates, n_A_ = animals (combined for all experiments/replicates), n_C_ = number of cells analyzed (combined for all experiments/replicates). For performing statistical tests on fractions, data were transformed by computing their square root before conducting Two tailed Student’s t-test (two groups) or ANOVA followed by Bonferroni correction (multiple groups). For individual measurements which were not normally distributed, either Mann-Whitney (2 groups) or Kruskal-Wallis test followed by Dunńs Correction (multiple groups) were performed. For comparing stacked graphs (groups showing proportion of a whole) Fischeŕs Exact test was used. P value <0.05 was considered to represent a statistically significant difference. P values are reported in figures only for those comparisons from which biological insights were drawn, thus the absence of a p-value does not necessarily indicate that a difference is non-significant.

To support statements about the lack of differences between experimental groups (where p-values are > 0.05) we computed the effect size of the differences using GPower (University of Düsseldorf) with these parameters: t-test tails = two, α = 0.05, power (1-β) = 0.8 and the sample size of both groups. Then, the smallest difference that would have been significant was calculated based on the effect size and the standard deviations and sample sizes of the experimental groups. These calculated smallest significant differences and the observed differences are reported in the Figure legends. If the calculated smallest significant difference was lower than differences reported for similar experiments in other studies, we concluded that our data sets had sufficient statistical power to detect similar effects. Thus, we consider that statements about the absence of differences are warranted in such cases.

## Acknowledgements

We thank Doris Weber and Janet Köhler for their contributions to fish care, Guoxin Sun for help with qRT-PCRs, Laura Kellerer for raising the *bmp7a*-/-mutants and the core facility “Light microscopy” of the Medical faculty of Ulm University for support with imaging. We thank Prof. Steffen Just and Dr. Bernd Gahr for help with neonatal mouse cardiomyocyte cultures. The Weidinger lab acknowledges funding by the Deutsche Forschungsgemeinschaft (DFG, German Research Foundation) via project ID 514204501, project ID 251293561 (Collaborative Research Centre [CRC] 1149), project ID 316249678 (CRC 1279), and project ID 450627322 (CRC 1506). M.D.V was also funded by an intramural grant of the Medical Faculty of Ulm University (“Baustein-Programm 3.2”). The research of K.S-K. is funded by the DFG within the project ID 450627322 (CRC 1506), by the Graduate Training Group GRK 1789, and by project ID 251293561 (CRC 1149). The D’Uva lab received funding from the European Union – NextGenerationEU through the Italian Ministry of University and Research under PNRR – M4C2-I1.3 Project PE_00000019 “HEAL ITALIA” to Gabriele Matteo D’Uva CUP J33C22002920006 and from the Italian Ministry of Health (RC-2024-2790614). L.W acknowledges funding by the DFG via project B3 in Research Training Group 2254 and project B03 in CRC 1506 (project ID 450627322). D. P. P., H. F. M. and M. I. are or were members of the International Graduate School in Molecular Medicine, Ulm University.

## Contributions

**Mohan Dalvoy Vasudevarao:** Conceptualization (lead); Investigation – most experiments using zebrafish, Formal analysis – zebrafish transcriptomics data; Visualization; Writing – review and editing. **Denise Posadas Pena**: Conceptualization (supporting); Investigation – experiments using Smad6 manipulation and *bmp7* mutants in zebrafish, Formal analysis – zebrafish transcriptomics data; Visualization; Writing – review and editing. **Michaela Ihle**: Investigation – DNA fiber spreading assays; Visualization; Writing – review and editing. **Chiara Bongiovanni:** Investigation – Mouse cardiomyocytes culture; Visualization; Writing – review and editing. **Pallab Maity:** Investigation – experiments in cultured fibroblasts; Visualization. **Simone Redaelli:** Methodology – creation and characterization of Smad6 expressing zebrafish transgenic lines; Writing – review and editing. **Kathrin Happ:** Methodology – creation and characterization of Smad6 expressing zebrafish transgenic lines. **Dominik Geissler:** Investigation – cardiomyocyte transcription data. **Hossein Falah Mohammadi**: Formal analysis – zebrafish transcriptomics data; Visualization. **Melanie Rall-Scharpf:** Resources – HSPCs. **Chi-Chung Wu:** Investigation – experiment using Noggin3 and Bmp2b overexpression transgenics; Writing – review and editing. **Karin Scharffetter-Kochanek:** Conceptualization (supporting); Funding Acquisition; Supervision. **Gabriele D’Uva:** Conceptualization (supporting); Funding Acquisition; Supervision; Writing – review and editing. **Arica Beisaw:** Resources – single cell data; Writing – review and editing. **Lisa Wiesmüller:** Conceptualization (supporting); Funding Acquisition; Supervision; Writing – review and editing. **Gilbert Weidinger:** Conceptualization (lead); Funding Acquisition; Methodology – creation of Smad6 expressing zebrafish transgenic lines; Project Administration; Supervision; Visualization; Writing – original draft; Writing – review and editing.

