## Supplementary Figures for "BMP signaling promotes heart regeneration via alleviation of replication stress"

### Supplement

**Table of contents**


Supplementary Figure 1 page 2

Supplementary Figure 2 page 3

Supplementary Figure 3 page 4

Supplementary Figure 4 page 5

Supplementary Figure 5 page 7

Supplementary Figure 6 page 9

Supplementary Figure 7 page 10

Supplementary Table 1 page 11

Supplementary Table 2 page 11

Supplementary Table 3 page 12

Supplementary Table 4 page 13

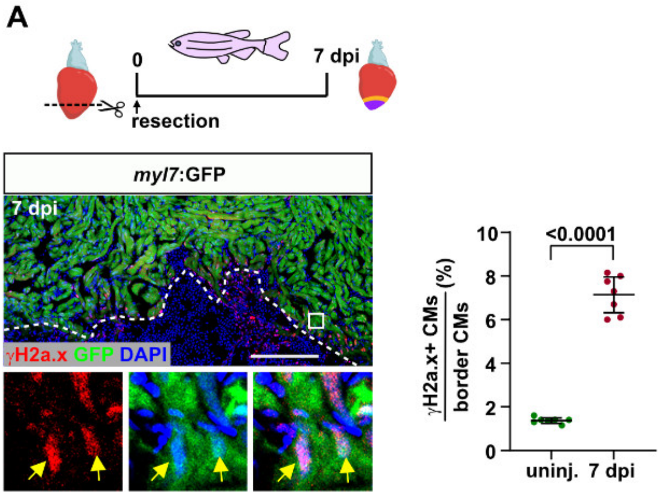


**Supplementary Figure 1**. *γH2a.x accumulates in wound border cardiomyocytes in hearts that regenerate after ventricular resection*

**(A)** Immunofluorescence on cryosections of *myl7*:GFP transgenic hearts reveals γH2a.x accumulation (arrows) in GFP+ CMs at the wound border at 7 days post ventricular resection (dpi). White box in the representative image indicates magnified region. Plot shows fraction of γH2a.x+ CMs out of all CMs within 150 µm of the wound border. Data points represent fraction in percent derived from individual hearts. Error bars, confidence interval (CI) 95%; Student´s t-test. $n_E$ = 1, $n_A$ = 7 uninjured, 7 resection, $n_C$ (analyzed CMs) = 5810 uninjured, 6120 resection. Scale bar, 150 µm.

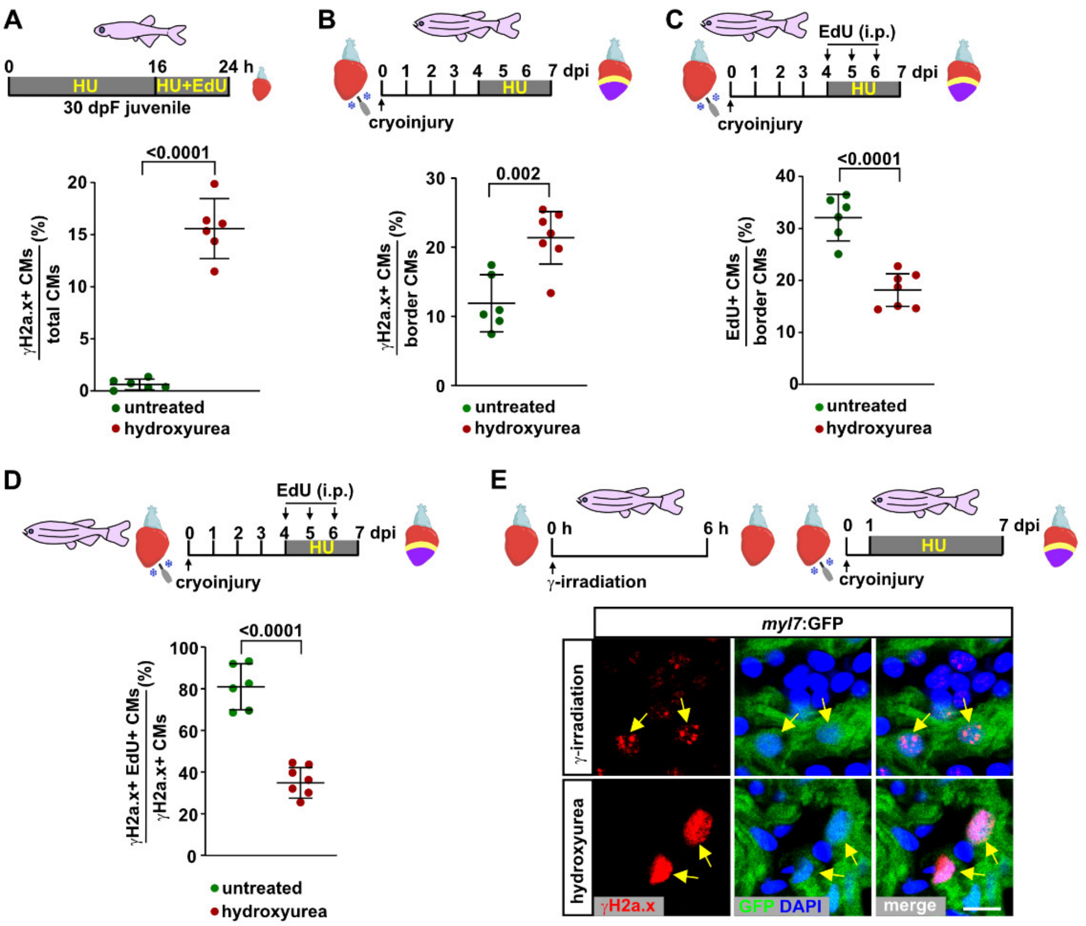


**Supplementary Figure 2**. *γH2a.x accumulation serves as reliable readout for replication stress in zebrafish cardiomyocytes*

(**A**) γH2a.x is hardly detectable in CMs of juvenile, uninjured fish, but induced by hydroxyurea treatment. Data are derived from the same fish as used for Figure 2B, which were also incubated with EdU as shown in the experimental scheme. Error bars, CI 95%; Student´s t-test. $n_E = 1$, $n_A = 6$ per condition, $n_C = 4560$ untreated, 3660 HU.

(**B**) Hydroxyurea treatment in adult injured fish further enhances the fraction of γH2a.x+ CMs within 150 µm of the wound border at 7 dpi. Error bars, CI 95%; Student´s t-test. $n_E = 1$, $n_A = 6$ untreated, 7 hydroxyurea, $n_C = 8360$ untreated, 7170 hydroxyurea.

(**C**) Hydroxyurea treatment in adult injured fish decreases the fraction of cycling wound border CMs labeled by daily i.p. injection of EdU from 4 to 6 dpi. Error bars, CI 95%; Student´s t-test. $n_E = 1$, $n_A = 6$ untreated, 7 hydroxyurea, $n_C = 8360$ untreated, 7170 hydroxyurea.

(**D**) Hydroxyurea treatment in adult injured fish decreases the fraction of γH2a.x+ wound border CMs at 7 dpi that are also EdU+ following EdU injection from 4 to 6 dpi. Error bars, CI 95%; Student´s t-test. $n_E = 1$, $n_A = 6$ untreated, 7 hydroxyurea, $n_C = 8360$ untreated, 7170 hydroxyurea.

(**E**) Immunofluorescence on cryosections of uninjured *myl7*:GFP transgenic hearts reveals γH2a.x accumulation in subnuclear foci in CMs 6 h after γ-irradiation (arrows). Intense pan-nuclear accumulation of γH2a.x is observed in the regenerating CMs of the wound border area at 7 dpi when fish are treated with hydroxyurea from 1 to 7 dpi (arrows). Representative image, not quantified. Scale bar, 10 µm.

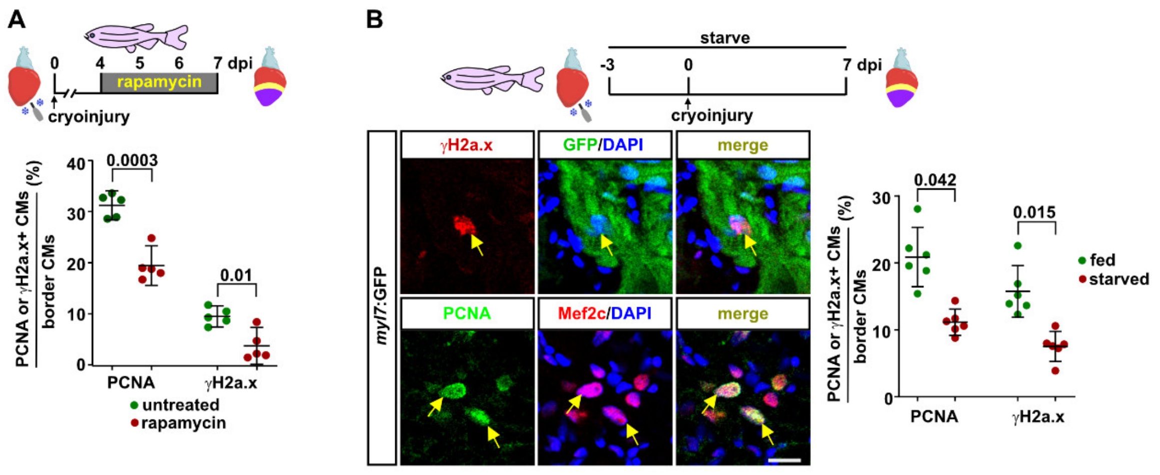


**Supplementary Figure 3**. *Reduction of cardiomyocyte proliferation is correlated with a decrease in γH2a.x accumulation*

**(A)** Incubation of cryoinjured fish with the mTOR inhibitor rapamycin for 3 days reduces the fraction of PCNA+ and of γH2a.x+ wound border CMs at 7 dpi. Error bars, CI 95%, Student´s t-test. $n_E$ = 1, $n_A$ = 5 per treatment, $n_C$ = 10540 untreated, 9700 rapamycin.

(**B**) Immunofluorescence on cryosections of *myl7*:GFP transgenic hearts reveals PCNA and γH2a.x accumulation in wound border CMs (identified by GFP or Mef2c staining) at 7 dpi in fish that are regularly fed. When fish are starved, the fraction of PCNA+ CMs as well as the fraction of γH2a.x+ CMs decreases. Error bars, CI 95%; Student´s t-test. $n_E$ = 1, $n_A$ = 6 per condition, $n_C$ = 12270 fed, 12720 starved. Scale bar, 10 μm.

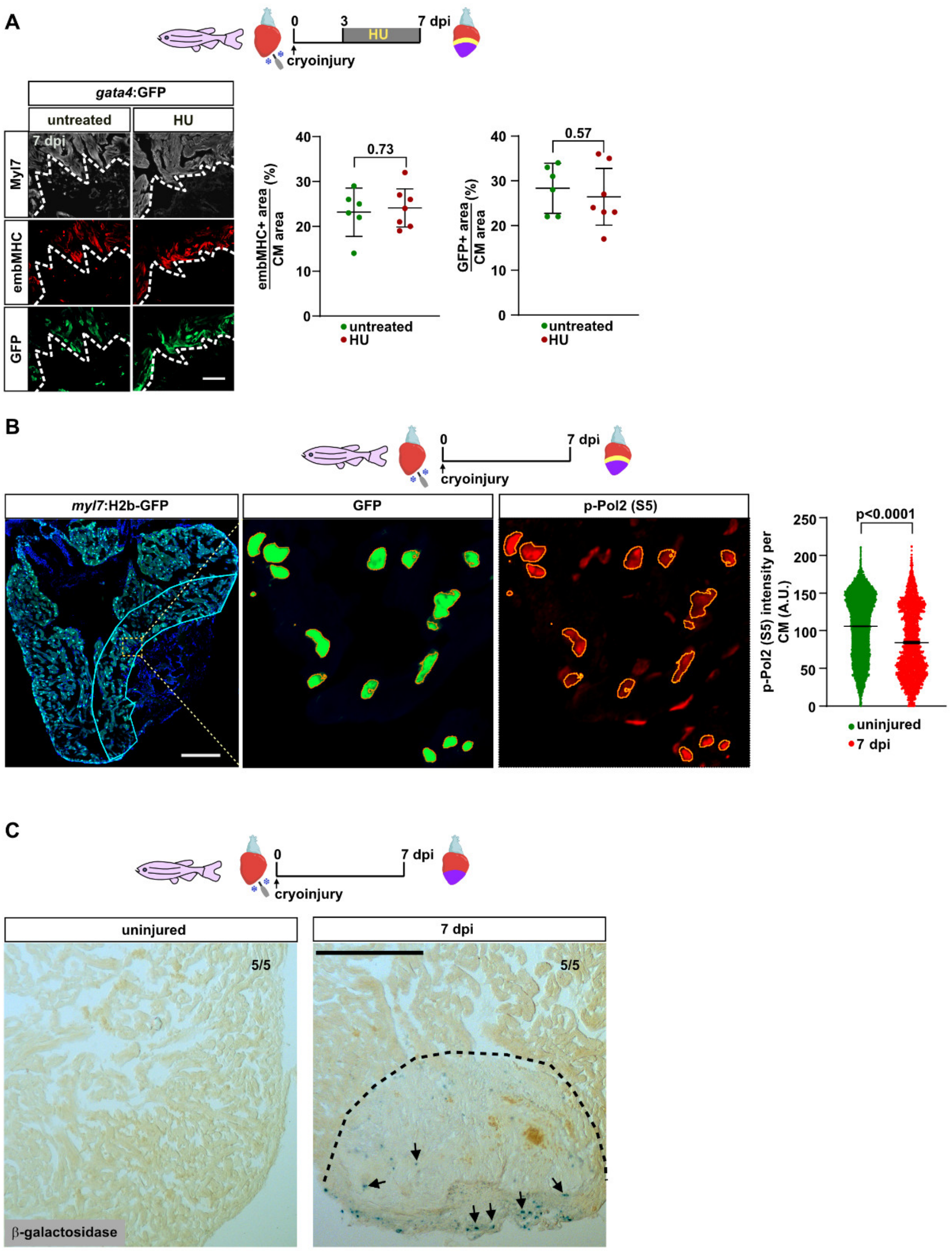


**Supplementary Figure 4**. *Replication stress does not effect CM dedifferentiation, is likely not caused by conflicts with transcription and does not induce CM senescence*

(**A**) Immunofluorescence on cryosections of *gata4*:GFP transgenic hearts reveals embryonic myosin heavy chain (embMHC) and GFP expression in Myl7+ CMs at the wound border at 7 dpi. The plots show the ventricular areas covered by anti-embMHC or anti-GFP staining relative to the 150 µm wound border zone area occupied by Myl7+. Replication stress induced by hydroxyurea treatment from 3 dpi until 7 dpi does not alter these readouts of CM dedifferentiation. Error bars, CI 95%; Student´s t-test. $n_E = 2$, $n_A = 6$ untreated, 7 HU.

For embMHC the observed relative difference between the untreated and HU treated groups is 7%, the calculated smallest significant difference 31%. For GFP the observed relative difference is 9%, the calculated smallest significant difference 33%. These calculated smallest significant differences are smaller than the effect sizes that we have observed previously after inhibition of Wnt signaling (172% and 87% respectively) {Bertozzi, 2022 #375}. We conclude that this experiment had enough power to detect biologically relevant effects. Since it did not, we conclude that HU treatment does not affect CM dedifferentiation.

(**B**) Wound border CMs display lower overall rates of transcription than remote CMs. Immunofluorescence on cryosections of *myl7*:H2b-GFP transgenic hearts reveals levels of p-Pol2 (S5), the active, elongating phosphorylated form of RNA Polymerase II, in CMs at the wound border at 7 dpi. Selections used to segment GFP+ nuclei are indicated in yellow, dotted box indicates magnified region. The intensity of p-Pol2 (S5) staining in nuclei is plotted in the wound border region of injured hearts at 7 dpi, and in a similarly sized myocardial region in uninjured hearts. Error bars, CI 95%; Student´s t-test. $n_E = 1$, $n_A = 5$ per condition, $n_S = 8$, $n_C = 18500$ uninjured, 4000 border CMs.

(**C**) Cryosections of uninjured and regenerating hearts at 7 dpi that had been stained for beta-galactosidase activity in the whole mount. At 7 dpi, sparse β-galactosidase-positive senescent cells are located within the wound and at higher density in the outermost layers of the wound, which most likely represent the epicardium that has already covered the wound (black arrows) No senescent cells can be detected in the wound border cardiomyocytes. Images are representative of 5 hearts per group.

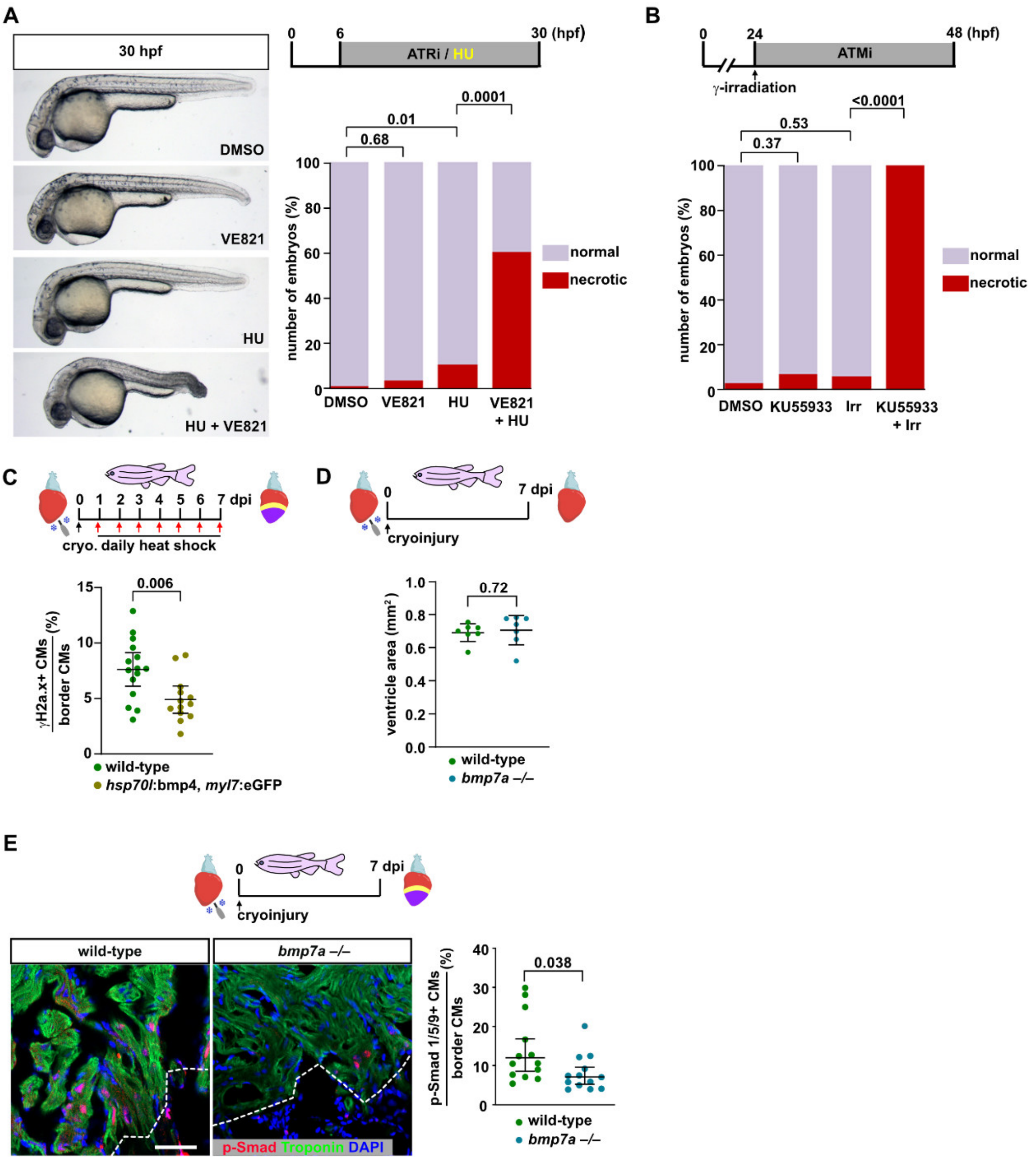


**Supplementary Figure 5**. *Specificity of ATM and ATR inhibitors in zebrafish, and further data on bmp4 overexpression and bmp7a mutant hearts.*

(**A**) The ATR inhibitor VE821 is not toxic to zebrafish embryos, but causes necrosis when applied in combination with hydroxyurea. Representative images of embryos at 30 hpf (hours post fertilization) show necrosis (darkened and deformed tissue) in the tail in HU + VE821 treated embryos. Plot shows the fraction of necrotic embryos. Fisher exact test. $n_E = 1$, $n_A = 30$ per treatment.

(**B**) The ATM inhibitor KU55933 is not toxic to zebrafish embryos, but causes necrosis when combined with γ-irradiation. Fisher exact test. $n_E = 1$, $n_A = 100$ per treatment.

(**C**) Bmp-GOF using *hsp70l*:bmp4 transgenic fish results in a decrease in the fraction of γH2a.x+ CMs in the wound border at 7 dpi compared to heat-shocked wild-type sibling hearts. Error bars, CI 95%; Student´s t-test. $n_E = 2$, $n_A = 15$ wild-type, 13 *hsp70l*:bmp4, $n_C = 7923$ wild-type, 7850 *hsp70l*:bmp4.

(**D**) Ventricles of *bmp7a –/–* mutant fish are similar in size to their wild-type siblings indicating that that hearts do not display obvious developmental defects. Data points represent total ventricle area derived from individual hearts at 7 dpi. Error bars, CI 95%; Student´s t-test. $n_E = 1$, $n_A = 7$ wild-type, 7 *bmp7a –/–*, $n_S$ = 8-12 per heart. The observed relative difference between wild-type and mutant groups is 2%, the calculated smallest significant difference 13%.

(**E**) *bmp7a –/–* mutant fish show reduced p-Smad1/5/9 in wound border CMs compared to wild type siblings at 7dpi. Error bars, CI 95%; Student´s t-test. $n_E = 2$, $n_A = 12$ wild-type, 13 *bmp7a –/–* $n_C = 4600$ wild-type, 6200 *bmp7a –/–*.

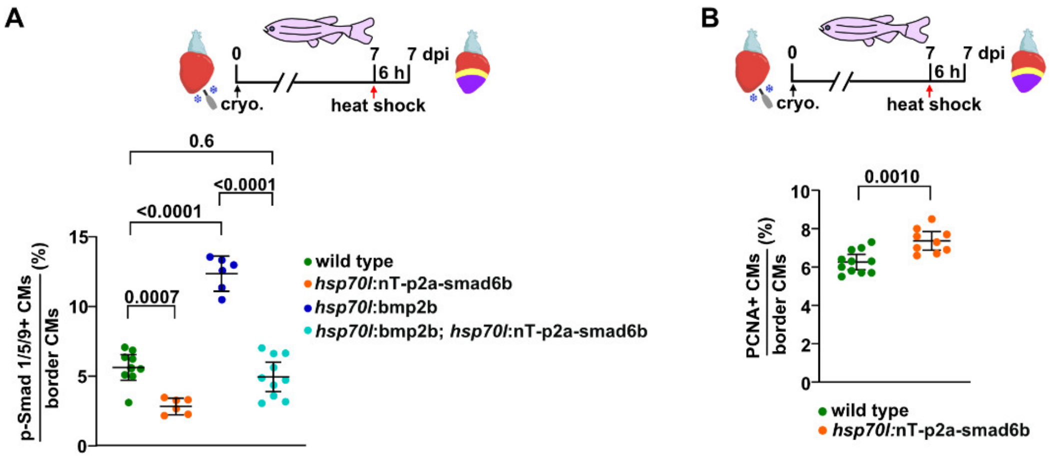


**Supplementary Figure 6.** *Smad6b and bmp2b overexpression cancel each other's ability to alter BMP signaling, and smad6b overexpression increases the fraction of PCNA+ CMs*

(**A**) Bmp-LOF using *hsp70l*:nT-p2a-smad6b reduces the fraction of p-Smad1/5/9+ wound border CMs 6 h after a single heat shock at 7 dpi, while *hsp70l*:bmp2b mediated Bmp-GOF increases it. Yet, in *hsp70l*:nT-p2a-smad6b; *hsp70l*:bmp2b double transgenics the fraction of p-Smad1/5/9+ CMs is comparable to that in heat-shocked wild-types, indicating that under these conditions the ability of the two transgenes to modulate Smad signaling cancel each other. Error bars, CI 95%, ANOVA with Bonferroni corrections. $n_E$ =2, $n_A$ = 9 wild-type, 6 *hsp70l:*nT-p2a-smad6b*,* 6 *hsp70l*:bmp2b*,* 10 *hsp70l:*nT-p2a-smad6b *x hsp70l*:bmp2b, $n_C$ = 12000 total CMs across all groups.

(**B**) In *hsp70l*:nT-p2a-smad6b transgenic hearts the fraction of PCNA+ CMs at the wound border is increased 6 h after a single heat-shock at 7 dpi. Error bars, CI 95%, Student´s t-test. $n_E$ =2, $n_A$ = 11 wild-type, 9 *hsp70l:*nT-p2a-smad6b*,* $n_C$ = 6800 wild-type, 6950 *hsp70l:*nT-p2a-smad6b.

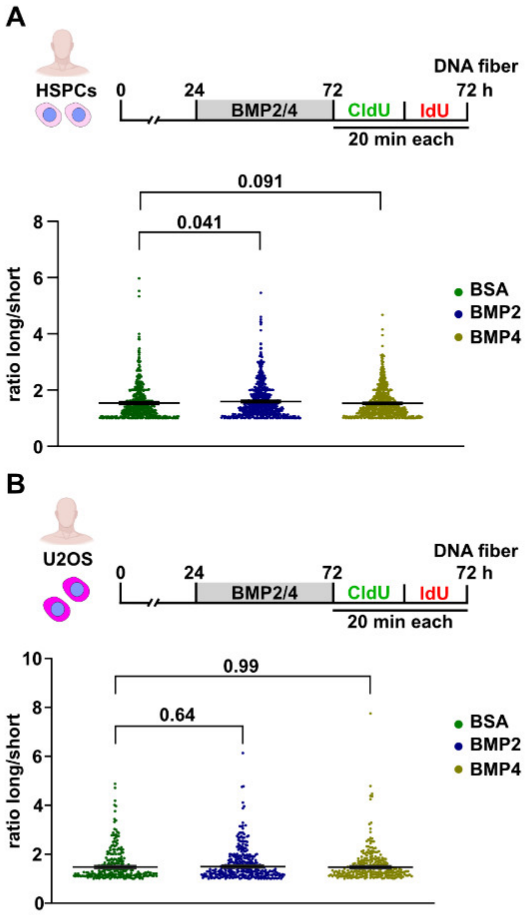


**Supplementary Figure 7.** *Pretreatment with BMP ligands does not affect replication fork stalling*

**(A)** Long vs short track ratios measured from DNA fibers containing both CldU and IdU tracks in Figure 8A indicate a minor, yet significant increase in fork asymmetry in hematopoietic stem and progenitor cells treated with BMP2, but not with BMP4 ligands. Data points represent the ratio of long to short tracks within individual CldU + IdU labeled fibers. Error bars, CI 95%, Kruskal-Wallis followed by Dunn´s correction test. $n_E = 3$, $n_{fibers} > 600$ per treatment.

**(B)** Long vs short track ratios measured from DNA fibers containing both CldU and IdU tracks in Figure 8B shows no change in fork asymmetry in U2OS cells treated with BMP2 or BMP4 ligands. Data points represent the ratio of long to short tracks within individual CldU + IdU labeled fibers. Error bars, CI 95%, Kruskal-Wallis followed by Dunn´s correction test. $n_E = 2$, $n_{fibers} > 250$ per treatment.

#### Supplementary Tables

#### Supplementary Table 1. Drugs, chemicals, ligands

| Name | Manufacturer | Identifier |
|---|---|---|
| Alfacalcidiol (vitamin D agonist) | Selleckchem | Cat# S1468 |
| Rapamycin | Selleckchem | Cat# S1039 |
| Hydroxyurea | Merck | Cat# H8627 |
| VE-821 (ATR Kinase Inhibitor) | Selleckchem | Cat# S8007 |
| KU55933 (ATM Kinase Inhibitor) | Selleckchem | Cat# S1092 |
| Nocodazole | Merck | Cat# M1404 |
| ClickTech EdU Cell Proliferation Kit 647 for IM | Baseclick | Cat# EdU647IM100+IV-S |
| Senescence β-Galactosidase Staining | Cell Signaling | Cat# 9860S |
| KASP Master mix | LGC Biosearch Technology | Cat# KBS-1050-101 |
| KASP Assay primer mix | LGC Biosearch Technology | Cat# KBS-2100-100 |
| 4',6-Diamidin-2-phenylindol -dihydrochlorid (DAPI) | Merck | Cat# 32670 |
| DMSO | Merck | Cat# D2650 |
| Recombinant Human BMP-2 | RandD systems | Cat# 355-BM-010 |
| Recombinant Human BMP-4 | RandD systems | Cat# 314-BP-010 |
| Recombinant Human BMP-7 | RandD systems | Cat# 354-BP-010 |
| SB202190 (p38 kinase inhibitor) | Sigma | Cat# S7067 |

#### Supplementary Table 2. Primers for RT-qPCR from zebrafish samples

| Gene | Identifier (ZFIN) | Sense primer | Antisense primer |
|---|---|---|---|
| *Rad51* | ZDB-GENE-040426-2286 | GTCATCACTAACCAGGTTGTAGC | ATCTCCCACTCCATCAGCATTAAT |
| *Rad54l* | ZDB-GENE-040426-968 | ATAGAGGAGAAGATCCTCCAGAGAC | AGTTGGAGAGATCACAGGTACAGTC |
| *Xrcc5* | ZDB-GENE-041008-108 | AGGAGCACTGAGTATTACACCAAGA | AATGACAGGAACTGATTTGCTTCT |
| *Gtf2h4* | ZDB-GENE-030131-6779 | ACACCCAGTAATGCTTAAACAGACC | AGAATCTCTTAACCTCGCTGTGTC |
| *Fen1* | ZDB-GENE-031112-11 | TAATTCAGTTCATGTGTGCTGAGAA | CGTAGCACTTGTCTTTTGTTTCTTT |
| *Rpa2* | ZDB-GENE-010131-3 | GTGCTAACATGATGCTAGTCAATGG | GATCTTCGTCGATGGTGGAGAAAAT |
| *Chaf1a* | ZDB-GENE-030131-5366 | CACAAACTCTTCTACCACACCCTC | CAAGGATGTGTTGATGTCCTGTTC |
| *Atm* | ZDB-GENE-040809-1 | AACAATGGAAGTTATGAGGAGTTCT | CACGCTCCGCCACTTTATTGAA |
| *Atr* | ZDB-GENE-070912-458 | TGTGAGGTCATACTAAGACTCATGA | TATTCCTGTTCTTGATCACTCCCTG |
| *Prkdc* | ZDB-GENE-030131-9008 | CTTGCTGCTCAACACTATGGATG | CCCTTATACTCCGGCATCTTCTC |
| *18s rRNA* | ZDB-RRNAG-180607-2 | CGCTATTGGAGCTGGAATTACC | GAAACGGCTACCACATCCAA |
| *Ubb* | ZDB-GENE-050411-10 | CTCAGATTAGAACCGACAGTCTTAGG | GCAACACAACATGAATAAAATAATGGGAAA |
| *Eef1a1l1* | ZDB-GENE-990415-52 | TTCCTCTTTCTGTTACCTGGCAA | CTTCTCGATGGTTCTCTTGTCGATT |

#### Supplementary Table 3. Primary antibodies

| Short name | Full name | Source | Identifier | Application |
|---|---|---|---|---|
| p-Smad1/5/9 | Rabbit monoclonal Phospho-Smad1 (Ser463/465)/ Smad5 (Ser463/465)/ Smad9 (Ser465/467) | Cell Signaling Technology | Cat# 13820 RRID:AB_2493181 | Immunostaining: Zebrafish cryosections |
| PCNA | Mouse monoclonal Proliferating Cell Nuclear Antigen | Dako | Cat# M0879 RRID:AB_2160651 | Immunostaining: Zebrafish cryosections |
| Mf20 | Mouse monoclonal MF20 (which detects sarcomeric Myosin heavy chain – MHC) | Developmental Studies Hybridoma Bank | Cat# MF20 RRID:AB_2147781 | Immunostaining: Zebrafish cryosections |
| Myl7 | Rabbit polyclonal Myl7 | GeneTex | Cat# GTX128346, RRID:AB_2885759 | Immunostaining: Zebrafish cryosections |
| Caspase-3 | Rabbit monoclonal Caspase 3 | BD Biosciences | Cat# 559565 RRID:AB_397274 | Immunostaining: Zebrafish cryosections |
| GFP | Chicken polyclonal GFP | Abcam | Cat# Ab13970 RRID:AB_2936447 | Immunostaining: Zebrafish cryosections |
| embMHC | Mouse monoclonal MYH7 | DSHB | Cat# N2.261 RRID:AB_531790 | Immunostaining: Zebrafish cryosections |
| pH3 | Mouse monoclonal Phospho-Histone 3 (Ser10) 6G3 | Cell Signaling | Cat# 9706S RRID:AB_331748 | Immunostaining: Zebrafish cryosections |
| p-Rpa32 (s33) | Rabbit polyclonal RPA32 Phospho (S33) | Bethyl Laboratories | Cat# A300-246A | Immunoblotting: Zebrafish ventricles |
| Gapdh | Rabbit monoclonal GAPDH | Cell Signaling | Cat# 2118S RRID:AB_561053 | Immunoblotting: Zebrafish ventricles |
| Mef2 | Rabbit polyclonal Mef2c | Santa Cruz Biotechnology | Cat# SC313 (discontinued) | Immunostaining: Zebrafish cryosections |
| Troponin | Cardiac troponin (CT3) | DSHB | Cat# CT3 | Immunostaining:Zebrafish cryosections |
| Troponin | Anti-cardiac troponin I | Abcam | Cat#ab47003 | Immunostaining: Primary neonatal mouse cardiomyocyte |
| γH2a.x | Rabbit polyclonal histone H2A.XS139ph (phospho Ser139) | Genetex | Cat# GTX127342 RRID:AB_2885642 | Immunostaining and immunoblotting: zebrafish cryosections and zebrafish ventricles |
| γH2a.x | Anti-phospho-Histone H2A.X (Ser139) Antibody, clone JBW301 | Millipore | Cat#05-636 | Immunostaining: Primary human neonatal dermal fibroblast |
| γH2a.x | Phospho-Histone H2A.X (Ser139) (20E3) | Cell Signaling | Cat#9718 | Immunostaining: Primary neonatal mouse cardiomyocyte |
| p-Pol2 (S5) | Anti-RNA polymerase II CTD repeat YSPTSPS (phospho S5) antibody | Abcam | ab5131 | Immunostaining: Zebrafish heart cryosections |
| BrdU, | Anti-BrdU detecting IdU (mouse) | BD Biosciences | Cat#347580 | DNA fiber spreading assays |
| BrdU | Anti-BrdU detecting CldU (rat, monoclonal, clone BU1/75 (ICR1)) | BioRad | Cat#OBT0030 | DNA fiber spreading assays |

#### Supplementary Table 4. Secondary antibodies

| Name | Source | Identifier | Application |
|---|---|---|---|
| Goat anti-Rabbit IgG (H+L) Highly Cross-Adsorbed Secondary Antibody, Alexa Fluor™ 488 | Invitrogen | Cat# A11034 RRID:AB_2576217 | Immunostaining |
| Goat anti-Rabbit IgG (H+L) Highly Cross-Adsorbed Secondary Antibody, Alexa Fluor™ 555 | Invitrogen | Cat# A21429 RRID:AB_2535850 | Immunostaining |
| Goat anti-Mouse IgG (H+L) Highly Cross-Adsorbed Secondary Antibody, Alexa Fluor™ 633 | Invitrogen | Cat# A21052 RRID:AB_2535719 | Immunostaining |
| Goat anti-Mouse IgG (H+L) Highly Cross-Adsorbed Secondary Antibody, Alexa Fluor 555 | Invitrogen | Cat# A21424 RRID:AB_141780 | Immunostaining |
| Goat anti-Rabbit IgG (H+L) Cross-Adsorbed Secondary Antibody, Alexa Fluor™ 633 | Invitrogen | Cat# A21070 RRID:AB_2535731 | Immunostaining |
| IRDye® 680RD Goat anti-Rabbit IgG Secondary Antibody | Licor | 926-68071 | Immunoblotting |
